# The translational landscape of SARS-CoV-2 and infected cells

**DOI:** 10.1101/2020.11.03.367516

**Authors:** Maritza Puray-Chavez, Nakyung Lee, Kasyap Tenneti, Yiqing Wang, Hung R. Vuong, Yating Liu, Amjad Horani, Tao Huang, Sean P. Gunsten, James B. Case, Wei Yang, Michael S. Diamond, Steven L. Brody, Joseph Dougherty, Sebla B. Kutluay

## Abstract

SARS-CoV-2 utilizes a number of strategies to modulate viral and host mRNA translation. Here, we used ribosome profiling in SARS-CoV-2 infected model cell lines and primary airway cells grown at the air-liquid interface to gain a deeper understanding of the translationally regulated events in response to virus replication. We find that SARS-CoV-2 mRNAs dominate the cellular mRNA pool but are not more efficiently translated than cellular mRNAs. SARS-CoV-2 utilized a highly efficient ribosomal frameshifting strategy in comparison to HIV-1, suggesting utilization of distinct structural elements. In the highly permissive cell models, although SARS-CoV-2 infection induced the transcriptional upregulation of numerous chemokines, cytokines and interferon stimulated genes, many of these mRNAs were not translated efficiently. Impact of SARS-CoV-2 on host mRNA translation was more subtle in primary cells, with marked transcriptional and translational upregulation of inflammatory and innate immune responses and downregulation of processes involved in ciliated cell function. Together, these data reveal the key role of mRNA translation in SARS-CoV-2 replication and highlight unique mechanisms for therapeutic development.

## INTRODUCTION

The Coronavirus (CoV) group encompasses of a number of single-stranded, positive-sense RNA viruses with unusually large genomes (27-32 kb), which infect a wide range of animal species, including humans (Masters, 2006, Weiss and Navas-Martin, 2005). Presently, SARS-CoV-2, the causative agent of the ongoing Coronavirus Disease-2019 (COVID-19) pandemic, continues to spread around the globe in part due to the emergence of viral variants with enhanced ability to transmit. Despite the high degree of protection with the SARS-CoV-2 vaccines, vaccine access remains limited globally. Furthermore there are limited options for antiviral or immunomodulatory treatment against SARS-CoV-2. A basic understanding of the replicative mechanisms of SARS-CoV-2 and associated host responses in relevant settings can foster the development of virus-specific therapies.

SARS-CoV-2 induced lung disease is thought to be in part due to manipulation of host type-I interferon (IFN) signaling (Sa Ribero et al., 2020). Compared to other CoVs such as SARS-CoV and Middle East respiratory syndrome (MERS), SARS-CoV-2 induces a poor or delayed IFN response in various experimental settings and *in vivo* (Blanco-Melo et al., 2020, Zhang et al., 2020, Lokugamage et al., 2020). Extensive characterization of SARS-CoV-2-encoded proteins within the past year has revealed multiple ways in which SARS-CoV-2 can post-transcriptionally manipulate host gene expression and induction of innate immune responses. For example SARS-CoV-2 NSP1, NSP6, NSP13, ORF3a, M, ORF7a and ORF7b inhibit STAT1/2 phosphorylation and ORF6 can inhibit STAT1 nuclear translocation (Sa Ribero et al., 2020, Lei et al., 2020, Konno et al., 2020, Miorin et al., 2020, Xia et al., 2020). NSP1 additionally binds to the mRNA entry channel of the 40S ribosomal subunit as well as non-translating 80S ribosomes to prevent binding of capped mRNA and thus inhibit the formation of the translation initiation complex (Schubert et al., 2020, Banerjee et al., 2020, Thoms et al., 2020, Lapointe et al., 2021). Furthermore, NSP16-mediated inhibition of alternative mRNA splicing has been implicated in suppression of innate immune responses (Banerjee et al., 2020). Under such inhibitory conditions SARS-CoV-2 mRNAs are thought to be efficiently translated owing to the structured elements within the 5’UTRs of viral mRNAs (Tidu et al., 2020, Banerjee et al., 2020, Finkel et al., 2021). On the other hand, the bulk of published research on SARS-CoV- and SARS-CoV-2-host interactions has relied on transcriptional profiling to study the immune response to infection (Blanco-Melo et al., 2020, Butler et al., 2020, Menachery et al., 2014, Mitchell et al., 2013, Wilk et al., 2020, Zhou et al., 2020). Such approaches may not fully capture the host immune response to infection, in the face of viral mechanisms that block host mRNA translation.

In addition to manipulation of host mRNA translation, SARS-CoV-2 utilizes programmed ribosomal frameshifting to successfully launch infection. The first two-thirds of the 5’ end of the SARS-CoV-2 genome is composed of two overlapping open reading frames (ORFs), ORF1a and ORF1b, which encode for two polyproteins, pp1a and pp1ab (Nakagawa et al., 2016). Pp1a is produced when translation of the genomic RNA terminates at the stop codon of ORF1a. Pp1ab is generated via a programmed - 1 ribosomal frameshift (PRF) that occurs at the overlap between ORF1a and ORF1b, permitting the elongating ribosomes to bypass the termination signal in ORF1a (Plant and Dinman, 2008). Following synthesis, pp1a and pp1ab are cleaved by viral proteases to generate 15-16 mature nonstructural proteins (NSPs) (Nakagawa et al., 2016). Many proteins encoded in ORF1b, are part of the replication complex, thus making the -1 PRF to generate pp1ab a critical translational event for SARS-CoV-2 replication. Frameshifting in coronaviruses is regulated by a highly conserved heptanucleotide slippery sequence (UUUAAAC) and an RNA pseudoknot structure a few nucleotides downstream (Plant and Dinman, 2008). The current models of PRF suggest that ribosomes stall upon encountering the pseudoknot (Plant et al., 2003, Korniy et al., 2019). This event presumably enhances the efficiency of ribosomal frameshifting by forcing the ribosomes to pause on the slippery sequence, which in turn promote the -1 slippage. Once the pseudoknot unwinds and resolves, the ribosomes can continue to translate the alternate ORF.

Another well-known frameshifting mechanisms in human viruses is employed by retroviruses through a stem-loop structure that regulates the expression of Gag/Gag-Pol transcripts (Jacks et al., 1988, Wilson et al., 1988). HIV-1 frameshifting is essential for maintenance of the ratio of Gag and Gag-Pol polyproteins as well as viral infectivity (Shehu-Xhilaga et al., 2001, Garcia-Miranda et al., 2016). While frameshifting is thought to be highly inefficient in HIV-1, with only 5-10% of ribosomes continuing into the Pol ORF (Baril et al., 2003, Dulude et al., 2006, Jacks et al., 1988), CoV frameshifting is thought to occur at a much higher efficiency (Irigoyen et al., 2016, Finkel et al., 2021, Finkel et al., 2020). Much of our understanding of viral frameshifting is based on reporter assays in transfected cells. However, to date, a comparison of SARS-CoV-2 and HIV-1 frameshifting efficiencies in infected cells has not been empirically assessed.

Here, we have conducted in-depth ribosome profiling studies to gain insight into the role of translational regulation in SARS-CoV-2 replication and the resulting host responses. We found that ribosome occupancy on viral mRNAs was temporally regulated and partly dependent on RNA abundance. In addition, ribosomes engaged with novel translation initiation sites (TIS) and other potential regulatory elements on SARS-CoV-2 RNAs. SARS-CoV-2 mRNAs quickly dominated the cellular mRNA pool but were not translated at a higher efficiency than cellular mRNAs overall. In addition, we found accumulation of ribosomes on the SARS-CoV-2 and HIV-1 frameshifting elements, but found that SARS-CoV-2 frameshifting was substantially more efficient than HIV-1. Remarkably, while numerous inflammatory chemokines, cytokines and ISGs were upregulated transcriptionally in SARS-CoV-2-infected Vero E6 cells, we found that many were not efficiently translated. Though we found that mRNAs encoding certain immune defense mediators were also less efficiently translated in primary airway cultures upon SARS-CoV-2 infection, repression of host mRNA translation in this physiologically relevant system was overall more modest. Taken together, our study defines the translational landscape of SARS-CoV-2-infected cells, revealing novel events that may promote viral replication and disarm host immune responses at the level of mRNA translation.

## RESULTS

### Ribosome profiling reveals key features of SARS-CoV-2 translational program

To study the relationship between transcriptionally and translationally regulated events at early and late phases of SARS-CoV-2 infection, Vero E6 cells infected at high multiplicity of infection (MOI) were monitored by RNA-seq and ribo-seq during the course of infection for 24 h (Fig. 1A). Viral antigen staining of infected cells revealed that the majority of the cells were infected by 12 hpi (**Fig. S1A**). Triplicate sequencing libraries (RNA-seq and ribo-seq) were generated and the mapping statistics are detailed in **Table S1, S2**. The quality of each sample and ribo-seq library was assessed as follows. First, despite the high degree of infection, RNA integrity was unaffected (**Fig. S1B**), suggesting that selection of poly-adenylated mRNAs for RNA-seq is unlikely to introduce a major 3’ bias. Second, the length of distribution of ribo-seq reads that mapped to cellular and viral transcriptomes were within the expected range of ribosome protected fragments (**Fig. S2A**) (Ingolia et al., 2012, Ingolia et al., 2009), although we noted that in one replicate experiment read lengths trended to be longer likely due to less extensive nuclease digestion (**Fig. S2A**). Third, irrespective of the differences in the average read-length distribution of independent experiments, the majority of ribo-seq reads mapped to coding sequences (CDS) and 5’ UTRs, with a clear reduction in the fraction of reads mapping to 3’UTRs when compared to RNA-seq experiments done in parallel (**Fig. S2B**). Finally, mapped ribosome-derived reads within the CDSs were enriched in fragments that align to the translated frame for cellular mRNAs (**Fig. S3A, B**). A similar outcome was observed for virally mapping reads except for one replicate where nuclease digestion was incomplete (**Fig. S4**).

**Figure 1.**
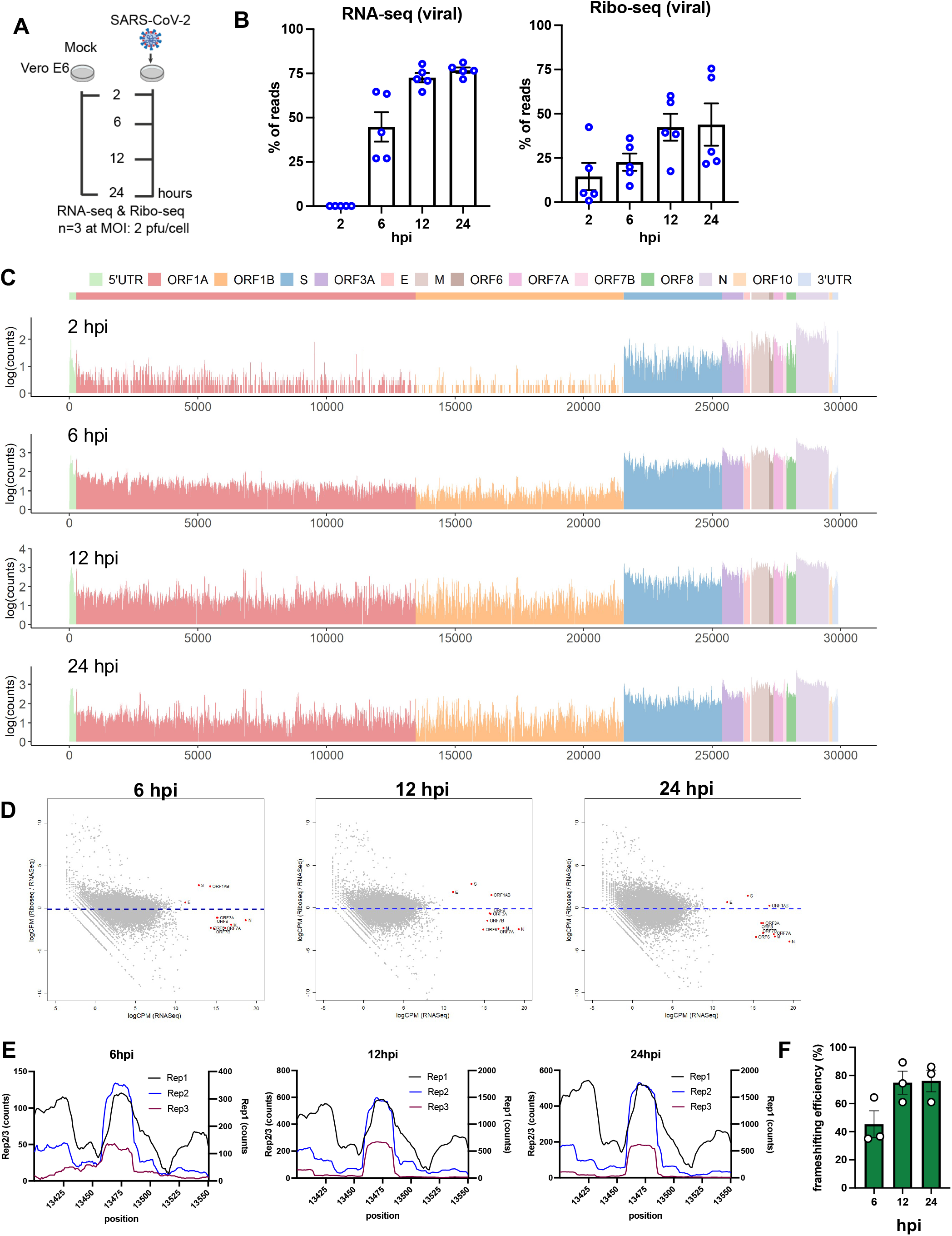
Ribo-seq reveals the translational program of SARS-CoV-2. (A) Schematic diagram of Ribo-seq and RNA-seq experiments conducted in this study. Vero E6 cells were infected at 2 pfu/cell and cells were processed for RNA-seq and Ribo-seq at 2, 6, 12 and 24 hpi. (B) Percentage of RNA-seq and Ribo-seq reads uniquely mapping to SARS-CoV-2 and cellular transcripts at the indicated time points post infection. Individual data points indicate independent biological replicates. (C) Ribo-seq counts along the viral genome across various time points. Schematic diagram of SARS2 genome features shown above is co-linear (also see **Table S3**). (D) Translation efficiency of viral (red) vs. host mRNAs (grey circles) are shown. (E) Ribo-seq read counts within the frameshifting site across three independent replicates is shown at 6, 12 and 24 hpi. (F) Data show the SARS-CoV-2 frameshifting efficiency as determined by comparing the average read densities between ORF1a and ORF1b regions across three independent replicates and various time points post-infection.

At 2 hpi, only a small fraction of mRNAs was derived from SARS-CoV-2 RNAs (Fig. 1B). At 6 hpi, a dramatic increase in vRNA levels was observed and by 12 hpi, nearly 80% of the total mRNA pool was viral (Fig. 1B). Viral RNAs were present abundantly in the ribosome bound pool as well and by 12 hpi ∼50% of the ribosome-protected fragments contained SARS-CoV-2 sequences (Fig. 1B). Plotting of RNA-seq reads on the SARS-CoV-2 genome demonstrated that N-derived sgRNAs were highly abundant throughout infection (**Fig. S5A, Table S3**), a finding consistent with previous RNA-seq studies (Kim et al., 2020, Huang et al., 2020). Ribosome density on SARS-CoV-2 mRNAs mirrored RNA abundance, with ribosomes enriched primarily on N-coding mRNAs (Fig. 1C**, S5B, Table S3**). The translational efficiency of viral mRNAs was not substantially different than the majority of cellular mRNAs with ORF1AB, S and E mRNAs translated at a modestly higher efficiency and the remainder of viral mRNAs translated at a lower efficiency than average, a pattern that did not vary with progression of infection (Fig. 1D). Thus, the high abundance of viral mRNAs, as opposed to a specific regulated mechanism, likely ensures the abundance of viral proteins, a finding consistent with other published studies (Finkel et al., 2021, Finkel et al., 2020).

In two replicate experiments, analysis of ribo-seq derived reads on viral RNAs at 2 hpi revealed the presence of a high occupancy site spanning nucleotides 27371-27457 (**Fig. S6A, Table S3**), which accounted for the majority of ribo-seq reads derived from viral RNAs at this time point. Further analysis of reads mapping to this region revealed an average read length distribution smaller than what is expected of ribosome protected fragments (**Fig. S6A**), suggesting that this peak is unlikely to be derived from RPFs. Ribosome occupancy on viral RNAs increased significantly by 6 hpi in all experiments, featuring S and downstream ORFs as the most frequently translated regions (Fig. 1C**, S5B**). Ribosome density noticeably increased also on the ORF1ab by 6hpi, but with lower read counts in ORF1b. Ribosome occupancy across viral RNAs increased further by 12 hpi and remained high during the remainder of infection (Fig. 1C**, S5B**). Ribosome footprints were non-uniform with numerous high and low frequency binding sites observed reproducibly across viral RNAs (Fig. 1C**, S5B**) with expected higher ribosome density within viral translation initiation sites (**Fig. S5B**).

Similar to other CoVs, SARS-CoV-2 frameshifting is thought to be mediated by a conserved heptanucleotide slippery sequence (UUUAAAC) and a RNA pseudoknot downstream from it spanning nucleotides 13408-13540 (**Fig. S6B**). A notable local increase in ribosome occupancy was observed surrounding the slippery site within the frameshifting element (Fig. 1E**, Fig. S6C, Table S3**), suggesting the possibility of steric hindrance by the FSE on translating ribosomes. Frameshifting was also evident in P-site analysis of the mapped reads with a notable shift from frame 0 to frame 2 (−1 frame), before and after the frameshifting site (**Fig. S6C).** Comparison of read density distribution between ORF1a and ORF1b indicated a relatively high efficiency of frameshifting ranging from %50 to %75 throughout the course of infection (Fig. 1F) in line with published reports for SARS-CoV-2 as well as other coronaviruses (Irigoyen et al., 2016, Dinan et al., 2019, Finkel et al., 2020).

We next tested whether SARS-CoV-2 can utilize alternative translation initiation, which is increasingly recognized as a key post-transcriptional regulatory mechanism (Kwan and Thompson, 2019, James and Smyth, 2018). To do so, ribo-seq experiments were performed in the presence of harringtonine, which results in the accumulation of ribosomes at translation initiation sites. In addition to enrichment of ribosomes at the canonical start codons, harringtonine treatment resulted in accumulation of ribosomes at alternative translation initiation sites during the course of infection, albeit at generally lower frequencies. For example, at 6 hpi, an internal noncanonical start codon ‘UUG’ within M ORF was utilized at ∼30% of the time, predicted to result in an out-of-frame peptide of 53 amino acids long (**Table S3, S4, Fig. S7**). An alternative translation initiation codon ‘AGG’ at 21868 nt appeared to be utilized within S at 6, 12 and 24 hpi, which would result in a short 18 amino acid peptide (**Table S3, S4, Fig. S7**). Finally alternative translation initiation sites were observed within M, resulting in an out-of-frame peptide and a truncated version of M (**Table S3, S4, Fig. S7**).

### HIV-1 frameshifting is regulated through a distinct mechanism compared with SARS-CoV-2

Analogous to SARS-CoV-2, HIV-1 also utilizes -1 ribosomal frameshifting, in this case for generation of the Gag-Pol polyprotein (Jacks et al., 1988, Wilson et al., 1988). HIV-1 frameshifting is regulated by a slippery sequence followed by a structured hairpin loop (Mouzakis et al., 2013, Staple and Butcher, 2005). To compare the frameshifting efficiency of SARS-CoV-2 to HIV-1, we next performed paired ribo-seq and RNA-seq experiments in HIV-1-infected CD4+ T-cells isolated from two independent donors (**Table S5 and S6**). Length distribution of ribo-seq derived reads that mapped to cellular and viral mRNAs were within the expected range of ribosome-protected fragments (**Fig. S8A**). In addition, ribo-seq reads that mapped to cellular mRNAs had a 3-nt periodicity in frame with annotated CDSs for varying read lengths (**Fig. S8B, S8C**) and were largely depleted of 3’ UTRs (**Fig. S8D**), suggesting that a large fraction of sequencing reads represent sequences derived from translating ribosomes.

We found that ribosome occupancy was high within the HIV-1 frameshifting element overlapping the slippery sequence (Fig.2A-C, **Table S7**) but dropped substantially 3’ to it and remained low throughout the Pol ORF (Fig. 2A, C, **Table S7**). Interestingly, another high occupancy site was observed immediately upstream of the slippery sequence within the FSE (Fig. 2C, **Table S7**). This suggests that ribosomes may pause and accumulate within the frameshifting site but only a small fraction of them continue translating into the Pol ORF, a finding that agrees with prior estimates of low (5-10%) HIV-1 frameshifting efficiency (Biswas et al., 2004, Dulude et al., 2006, Shehu-Xhilaga et al., 2001, Baril et al., 2003, Jacks et al., 1988). Thus, we conclude that programmed ribosome frameshifting is regulated through distinct mechanisms and possibly structures between HIV-1 and SARS-CoV-2. Interestingly, while the translation efficiency of *Env*, *Nef* and *Pol* mRNAs were similar to cellular mRNAs, the translation efficiency of the Gag ORF was significantly higher (Fig. 2D), and in part may be ascribed to the unique structural elements present within the 5’UTR of Gag-coding mRNAs (Kharytonchyk et al., 2016).

**Figure 2.**
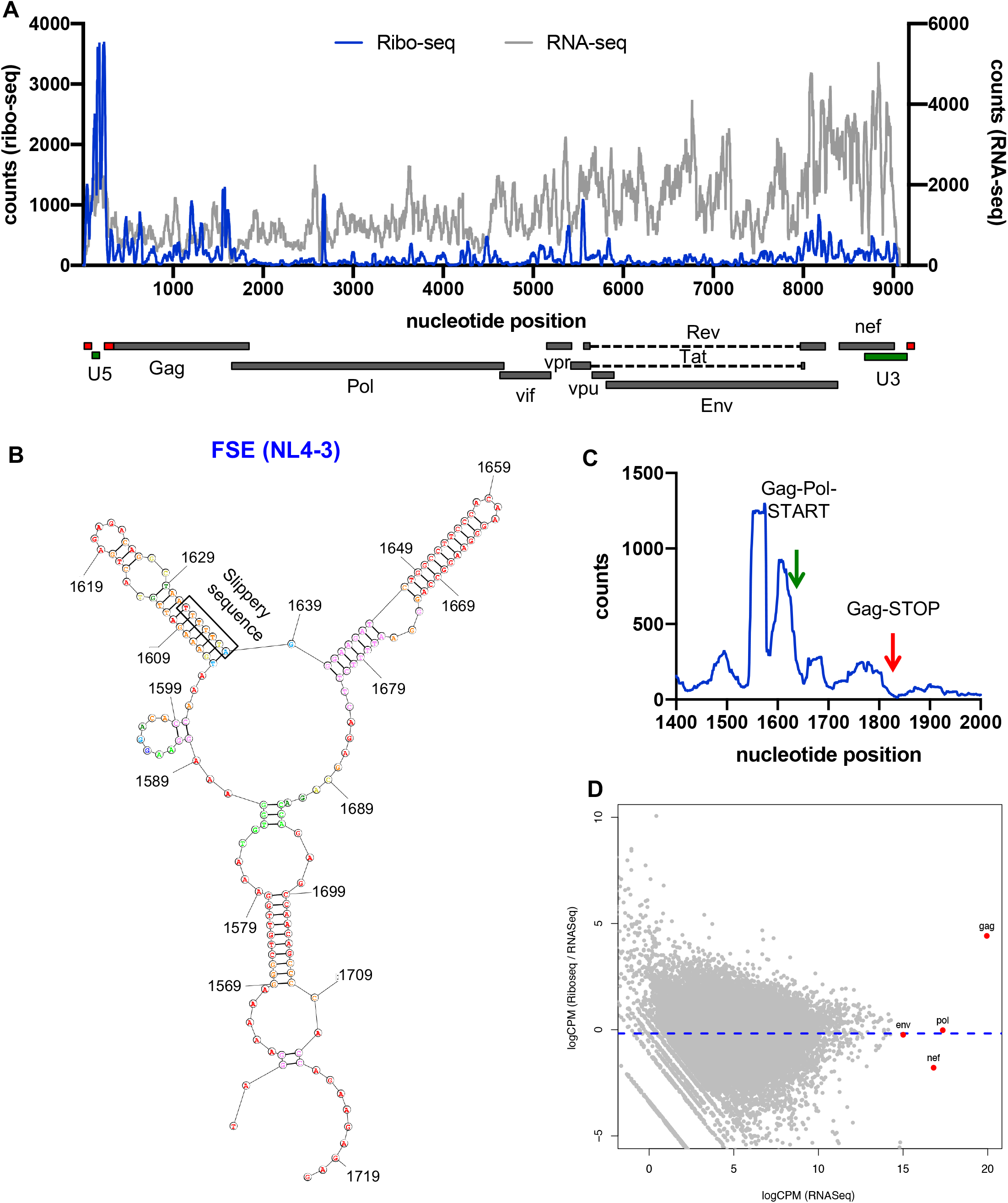
Ribo-seq in HIV-1-infected cells reveals inefficient ribosomal frameshifting. Primary CD4+ T-cells were infected with HIV-1_NL4-3_/VSV-G at an MOI of 2 and infected cells processed for RNA-seq and Ribo-seq at 24 hpi. (A) Ribo-seq and RNA-seq reads (counts) mapping to the HIV-1 genome are shown. Schematic diagram of HIV-1 genome features shown below is co-linear (also see **Table S7**). (B) Secondary structure prediction of the HIV-1 ribosome frameshifting element is shown. (C) Ribosome occupancy within the frameshifting site is illustrated. (D) Translation efficiency of viral (red) vs. host mRNAs (grey circles) is shown.

### Ribo-seq in primary HBECs reveal a similar SARS-CoV-2 translational program

SARS-CoV-2 primarily infects ciliated and type 2 pneumocyte cells in the human lung (Schaefer et al., 2020). Differentiated primary airway epithelial cells grown at the air-liquid interface (ALI) represent one of the most physiologically relevant models to study SARS-CoV-2 infection in culture. To corroborate the above findings from Vero E6 cells, we performed ribo-seq studies in SARS-CoV-2-infected primary human bronchial epithelial cells (HBEC) grown at the air-liquid interface (ALI). Cells inoculated at an MOI of 1 were processed for RNA-seq and ribo-seq at 4, 24, 48, 72 and 96 hpi (Fig. 3A). In contrast to the highly permissive Vero E6 cells, the progression of infection in HBECs was relatively slow and a small percentage of the cells were infected by 4 and 24 hpi (not shown). SARS-CoV-2 spread was visible by 48 hpi and a large fraction of ciliated cells expressing ACE2 were infected by 96 hpi (**Fig. S9A, 9B**). In agreement, the amount of newly synthesized viral RNAs was low at 4 hpi, but by 48 hpi approximately 20% of reads were of viral origin and did not increase further at 72 and 96 hpi (Fig. 3B, **Table S8, Table S9**). Of the relatively small number of RNA-seq-derived reads that mapped to the viral RNAs at 4 hpi, the majority were derived from subgenomic viral mRNAs coding for N and to a lesser extent from upstream ORFs including M, ORF6, ORF7 and ORF8 (**Fig. S10, Table S10**). Subgenomic viral mRNAs coding for N remained as the predominant species at later time points with notable increases at the expression level of upstream genes (**Fig. S10, Table S10**).

**Figure 3.**
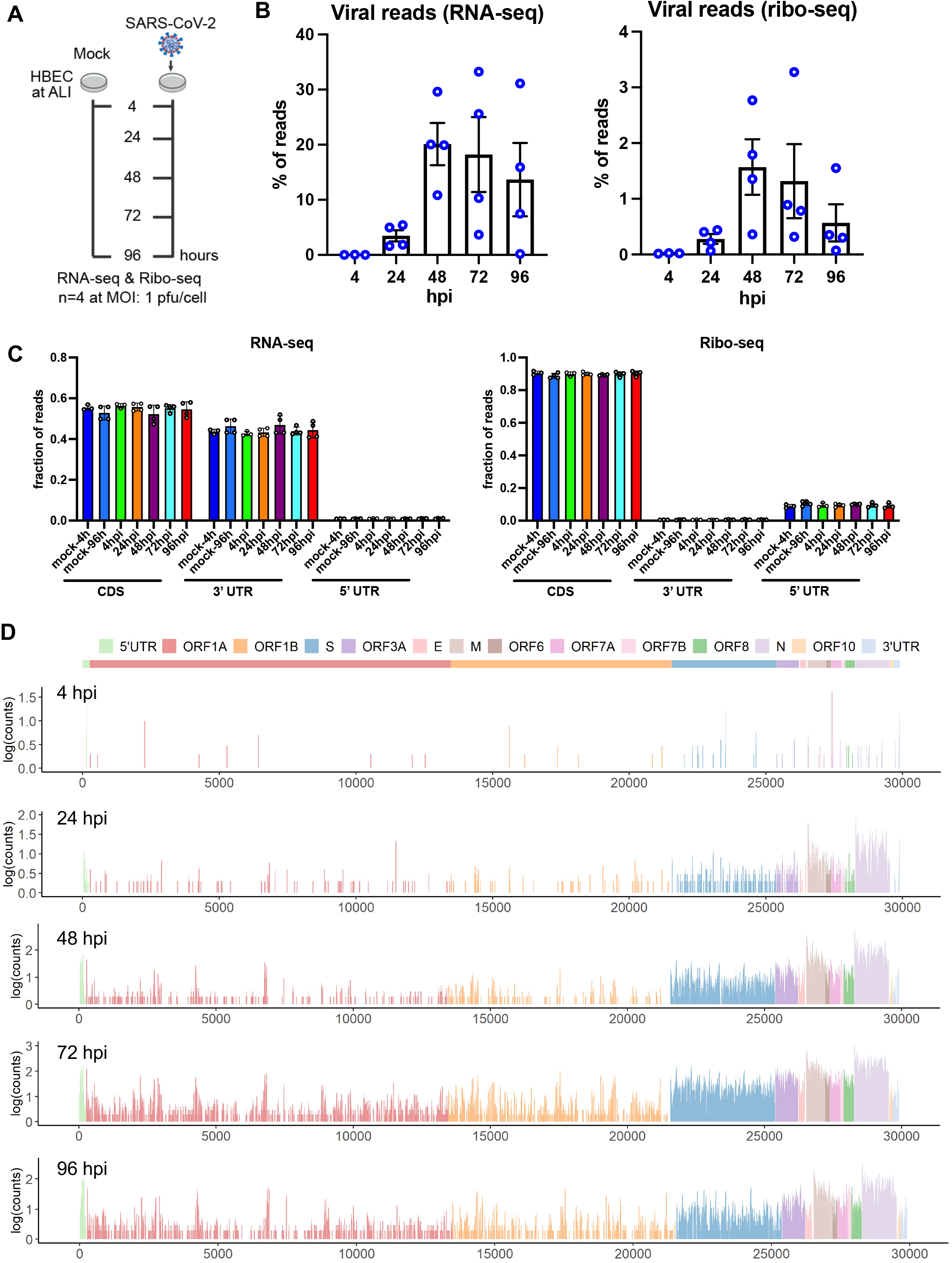
Ribo-seq in primary airway cells reveals a more restrictive translational program for SARS-CoV-2. (A) Schematic diagram of Ribo-seq and RNA-seq experiments conducted in this study. HBECs grown at ALI were infected at 1 pfu/cell and cells were processed for RNA-seq and Ribo-seq at 4, 24, 48, 72 and 96 hpi. (B) Percentage of RNA-seq and Ribo-seq reads uniquely mapping to SARS-CoV-2 and cellular transcripts at the indicated time points post infection. Individual data points indicate independent biological replicates. (C) Fraction of RNA-seq and ribo-seq-derived reads mapping to 5’UTRs, CDSs and 3’UTRs is shown. (D) Ribo-seq counts along the viral genome across various time points. Schematic diagram of SARS2 genome features shown above is co-linear (**also see Table S10**).

Quality of ribo-seq libraries was assessed as follows. First, as with previous experiments, RNA integrity was high despite widespread infection at 96 hpi (**Fig. S11A**). Second, length distribution of ribo-seq reads mapping to cellular and viral mRNAs matched the size expected from ribosome-protected fragments (**Fig. S11B**). Third, reads mapping to the 3’UTRs were depleted in ribo-seq libraries (Fig. 3C). Fourth, ribo-seq libraries were enriched in fragments that align to the translated frame and had a dominant frame with a 3-nt periodicity across varying read lengths for both cellular and virally mapping reads (**Fig. S12, S13**).

In contrast to Vero E6 cells, viral RNAs constituted only a small fraction of ribo-seq-derived RNAs (Fig. 3B) suggesting a significantly more restrictive translational environment overall for SARS-CoV-2 in ALI cultures. Ribosomes bound by viral RNAs were readily detected at 24, 48, and 72 hpi, but not at 4 hpi, with N and M ORFs being the most frequently translated (Fig. 3C). Overall translation efficiency of SARS-CoV-2 mRNAs was by and large proportional to the abundance of sgRNAs and proceeded in a similar cascade in the primary HBECs as well as in the Vero E6 cells. Due to the relatively low read coverage across ORF1ab, we did not assess frameshifting efficiency in this experimental setting.

### Inflammatory and innate immune mRNAs are inefficiently translated in SARS-CoV-2 infected cells

Parallel analysis of ribo-seq and RNA-seq data sets provides a powerful tool to analyze translational level changes in response to SARS-CoV-2 infection. Paired RNA-seq and ribo-seq data obtained from three independent experiments were analyzed for differential gene expression patterns in Vero E6 cells. Principal component analysis (PCA) showed samples separated well based on time post-infection despite a level of variability in particular at the 6 hpi time point (**Fig. S14A-C**).

Hierarchical consensus clustering of the 1,018 differentially expressed genes (DEGs) (|logFC|>2 and q<0.05) from RNA-seq generated 5 temporally resolved clusters (Fig. 4A, **S15A, Table S11**). As early as 2 hpi, we found transcriptional upregulation of transcription factors involved in cell cycle regulation and induction of inflammation (i.e.NR4A3 and EGR3) (Fig. 4A, **S16A, Table S11).** Numerous chemokine ligands (CXCL1, CXCL3, CXCL11, cluster 1) as well as IFN-α/β signaling and downstream ISGs significantly increased at 6 and 12 hpi (cluster 3; Fig 4A, **S15A, S16A, Table S11**). Induction of inflammatory and innate immune pathways were confirmed by gene set enrichment analysis (GSEA) of genes at each time point (Fig. 4B, **Table S11**). Another cluster (cluster 2) of upregulated genes were composed of genes involved in mRNA processing and mRNA translation (Fig. 4A, 4B, **Table S11**). Though numerous genes in clusters 4 and 5 were downregulated in all replicate experiments, we did not observe the specific enrichment of a pathway in this set of DEGs.

**Figure 4.**
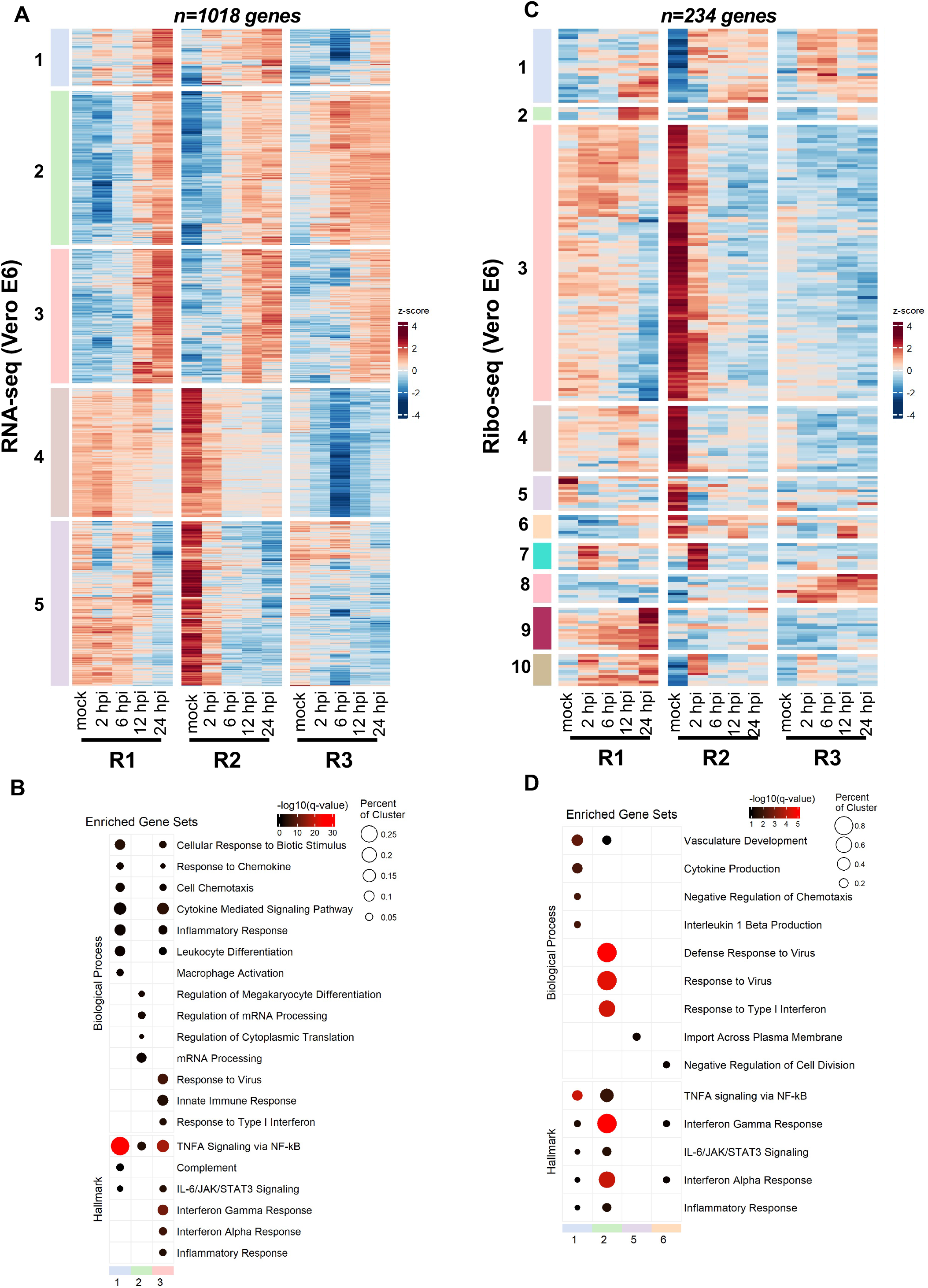
SARS-CoV-2 infection induces translational repression of innate immune genes. Vero E6 cells infected at 2 pfu/cell as detailed in Fig.1 were analyzed for differential expression of host genes by RNA-seq (A, B) and ribo-seq (C, D). (A, C) Hierarchical clustering of differentially expressed genes (DEGs) after infection are shown. Genes were filtered for an absolute log_2_ fold change >2 and adjusted q-value <0.05 at any time point. (B, D) Hypergeometric enrichment analysis from two different databases for each individual cluster in 4A and 4C (Hallmark, Gene Ontology). Color represents significance (q-value); size indicates the percentage of the cluster represented in the pathway. (Also see **Table S11, S12**).

Remarkably, the majority of these transcript level changes were not apparent in Ribo-seq data (Fig. 4C, 4D, **S15B, S16B, Table S11**). Only 234 genes were found to be differentially regulated in response to SARS-CoV-2 infection forming 10 temporally-resolved clusters (Fig. 4C, **S15B, Table S11**). Many of these clusters were smaller in size and the degree of differential expression varied in clusters 7-10 between replicate experiments (Fig. 4C, **S15B, Table S11**). Notwithstanding, cluster 2 (ribo-seq), which was enriched for type-I IFN response pathway was substantially smaller in size and consisted of only a few ISGs (i.e. IFIT1, IFIT2, IFIT3 and CXCL10) (Fig. 4C, **S15B, S16B Table S11**). In contrast, we found that another innate immune modulator, IL11, was significantly upregulated translationally but not transcriptionally at 2hpi (**Fig. S16B, Table S11**). Clusters 3 and 4 consisted of genes that were downregulated significantly but were not significantly enriched for a particular pathway (**Table S11**). Together, these findings suggest that immune response genes are translationally repressed, and their expression significantly delayed in infected cells.

Many of these findings held consistent for the RNA-seq and ribo-seq experiments performed on Vero E6 cells infected at a low MOI (**Table S12, S13**). For example, transcription factors ATF3 and EGR1, key regulators of inflammatory responses, were upregulated at 24 hpi alongside with numerous chemokine ligands (i.e. CXCL1, CXCL8, CXCL10) and interleukin 6 (**Fig. S17A, Table S14**). We also noted the upregulation of numerous ISGs (i.e. IFIT1, IFIT2, IFIT3) as well as IFN-lambda at 24 hpi (**Fig. S17A, Table S14**). The 48 hpi timepoint was marked by upregulation of genes involved in cell cycle regulation and apoptosis (i.e. FOS, NR4A3), as well as inflammatory cytokines such as IL-31 and ISGs including OASL (**Fig. S17B, Table S14**). In line with our above observations, the great majority of the transcriptionally upregulated genes were not translationally upregulated at 24 hpi (**Fig. S17B, Table S14**) and 48 hpi (**Fig.S17B, Table S14**). Interleukins IL11 and IL1A stood out as immune-related genes that were translationally upregulated at 24 and 48 hpi, respectively (**Fig. S17B, Table S14**).

Paired RNA-seq and ribo-seq data obtained from four independent infections of HBECs grown at ALI (from two independent donors) were analyzed for differential gene expression similarly. Principal component analysis (PCA) showed samples separated well based on time post-infection as well as donor samples (**Fig. S18A**). Furthermore, the level of gene level biological variability was within a reasonable range for both RNA-seq and ribo-seq libraries (**Fig. S18B, C**). SARS-CoV-2 infection induced differential expression of 2727 and 1208 genes in RNA-seq and ribo-seq experiments, respectively (**Table S15, S16**). As expected from the low level of infection at 4 hpi, relatively few genes were differentially regulated at this time point for both RNA-seq and ribo-seq data sets (**Table S15, S16**). Transcriptionally upregulated genes formed six temporally resolved clusters (Fig. 5A, **S19A, S20A, Table S15**). Cluster 2, which contained the largest number of upregulated DEGs, was significantly enriched in genes in the type I/III IFN pathway and inflammatory responses (Fig. 5B, **Table S15**). Clusters 4 and 5 were composed of genes that were downregulated at later stages of infection (Fig. 5A, **S19A, S20A, Table S15**). GSEA revealed that many of these genes are involved in cilium organization and movement (Fig. 5B, **Table S15**), demonstrating the impact of SARS-CoV-2-induced remodeling and/or killing of the ciliated cells in the airway cultures. In contrast to Vero E6 cells, the majority of these patterns were maintained in ribo-seq experiments. The 1208 DEGs derived from the ribo-seq experiments formed 5 clusters, with clusters 1-3 consisting of translationally upregulated genes (Fig. 5C, **S19B, 20B, Table S16**). While clusters 1 and 3 were not enriched in genes in a specific pathway, we found that Cluster 2 was significantly enriched in genes in the IFN and inflammatory response pathways (Fig. 5D, **Table S16**). Similar to RNA-seq data, GSEA of downregulated DEGs in cluster 5 revealed enrichment of genes involved in cilium organization and motility (Fig. 5D, **Table S16**).

**Figure 5.**
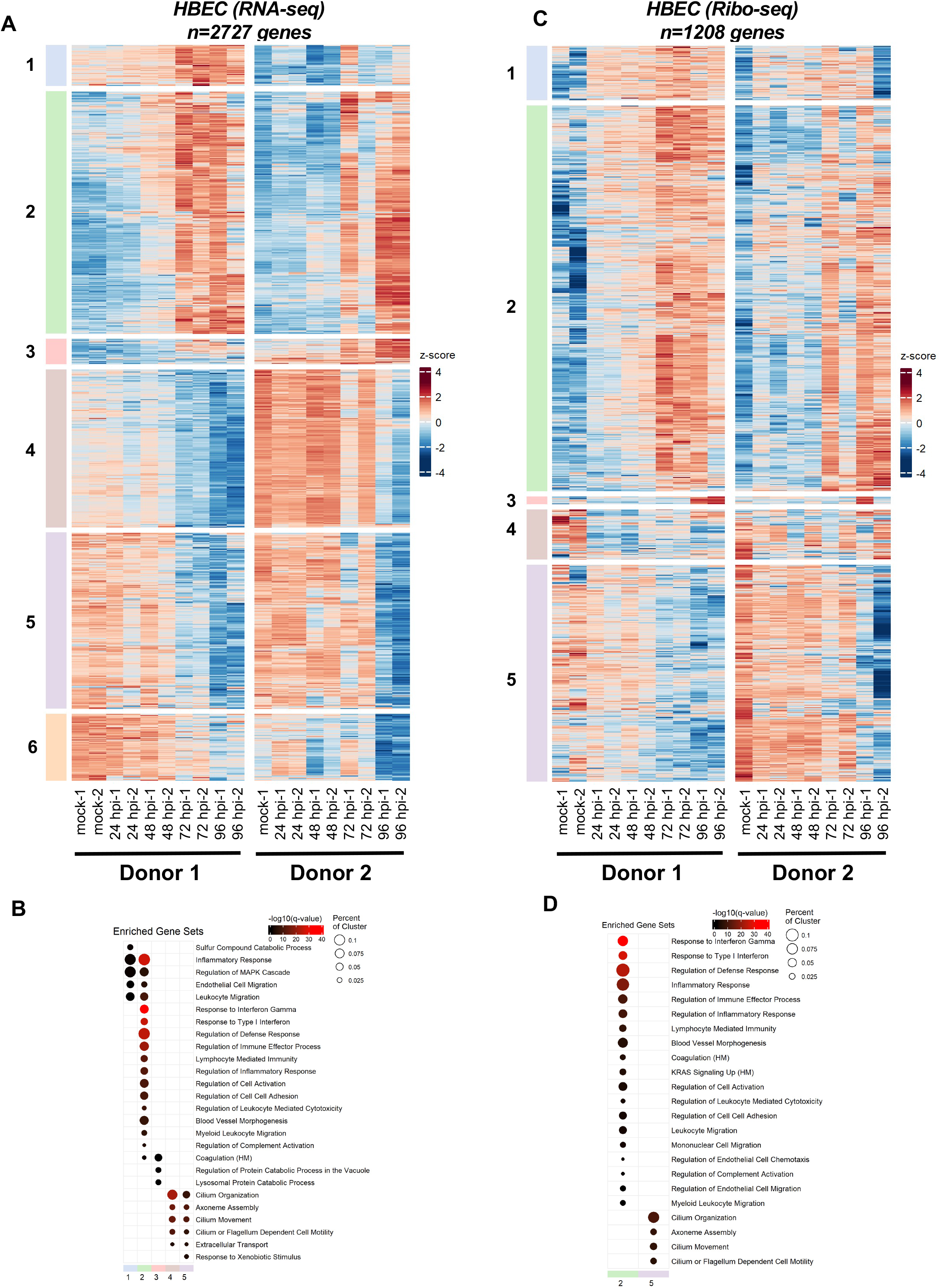
SARS-CoV-2-induced changes in primary airway cells. Primary human bronchial epithelial cells grown at air-liquid interface were infected at 1 pfu/cell as detailed in Fig.3 were analyzed for differential expression of host genes by RNA-seq (A, B) and ribo-seq (C, D). (A, C) Hierarchical clustering of differentially expressed genes (DEGs) after infection are shown. Genes were filtered for an absolute log_2_ fold change >2 and adjusted q-value <0.05 at any time point. (B, D) Hypergeometric enrichment analysis from two different databases for each individual cluster in 4A and 4C (Hallmark, Gene Ontology). Color represents significance (q-value); size indicates the percentage of the cluster represented in the pathway (see also **Table S15, S16**).

### Comparison of SARS-CoV-2 and host mRNA translation efficiencies

We next compared the translational efficiency of cellular host response genes in Vero E6 and primary HBEC cultures. In Vero E6 cells the translation efficiency of various immune modulatory genes was substantially lower than other cellular mRNAs, most evident at 12 and 24 hpi which marks the accumulation of viral proteins (Fig. 6A-D, **Table S17**). In contrast, in HBEC-ALI cultures, the translation efficiency of mRNAs encoding ISGs and inflammatory genes did not appear to be significantly lower than other cellular mRNAs (Fig. 6E-H, **Table S18**). Notable exceptions included CXCL9 and IFN-B, which were substantially upregulated at 48hpi at the transcript level but had lower translation efficiencies (Fig. 6F, **Table S18**).

**Figure 6.**
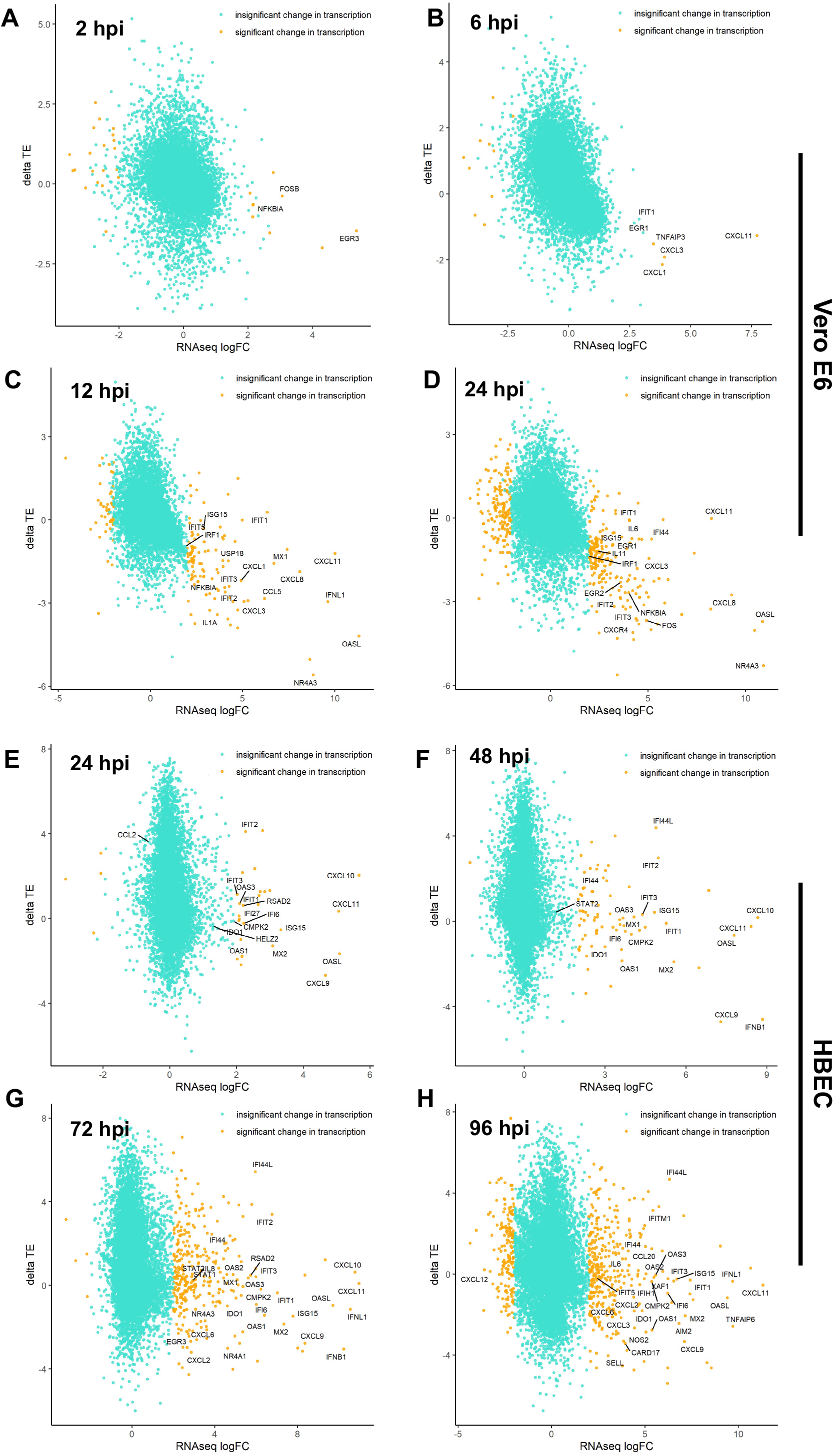
SARS-CoV-2-induces translational repression of innate immune genes. Changes in the translational efficiency of genes that were differentially transcribed in response to SARS-CoV-2 infection are shown for Vero E6 cells (A-D) and HBECs (E-H) at the indicated time points post-infection (see also **Table S17, 18**).

Numerous viral proteins have been implied in modulation of type-I IFN responses and we next tested the direct impact of some of these factors in suppression of ISG expression. To this end, cells transfected with Nsp1, Nsp7, ORF3a and ORF6 expression constructs were stimulated with IFN-α and induction of ISGs assessed by immunoblotting and Q-RT-PCR. We found that Nsp1 overexpression significantly reduced IFN-α-mediated phosphorylation of STAT1 (Fig. 7A), whereas other viral proteins had no impact on STAT1 levels or phosphorylation. Nsp1-mediated suppression of STAT1 phosphorylation was accompanied by a significant reduction of ISG upregulation at both protein and RNA level (Fig. 7A-C). While ORF3a and ORF6 did not affect STAT1 phosphorylation, they both reduced steady state ISG expression (Fig. 7A), yet the impact on ISG protein levels was relatively modest (Fig. 7B). These findings suggested that the observed translational repression of innate immune modulators in SARS-CoV-2-infected cells is likely due to the actions of multiple viral proteins and possibly due virus-induced changes and stress in heavily infected cells.

**Figure 7.**
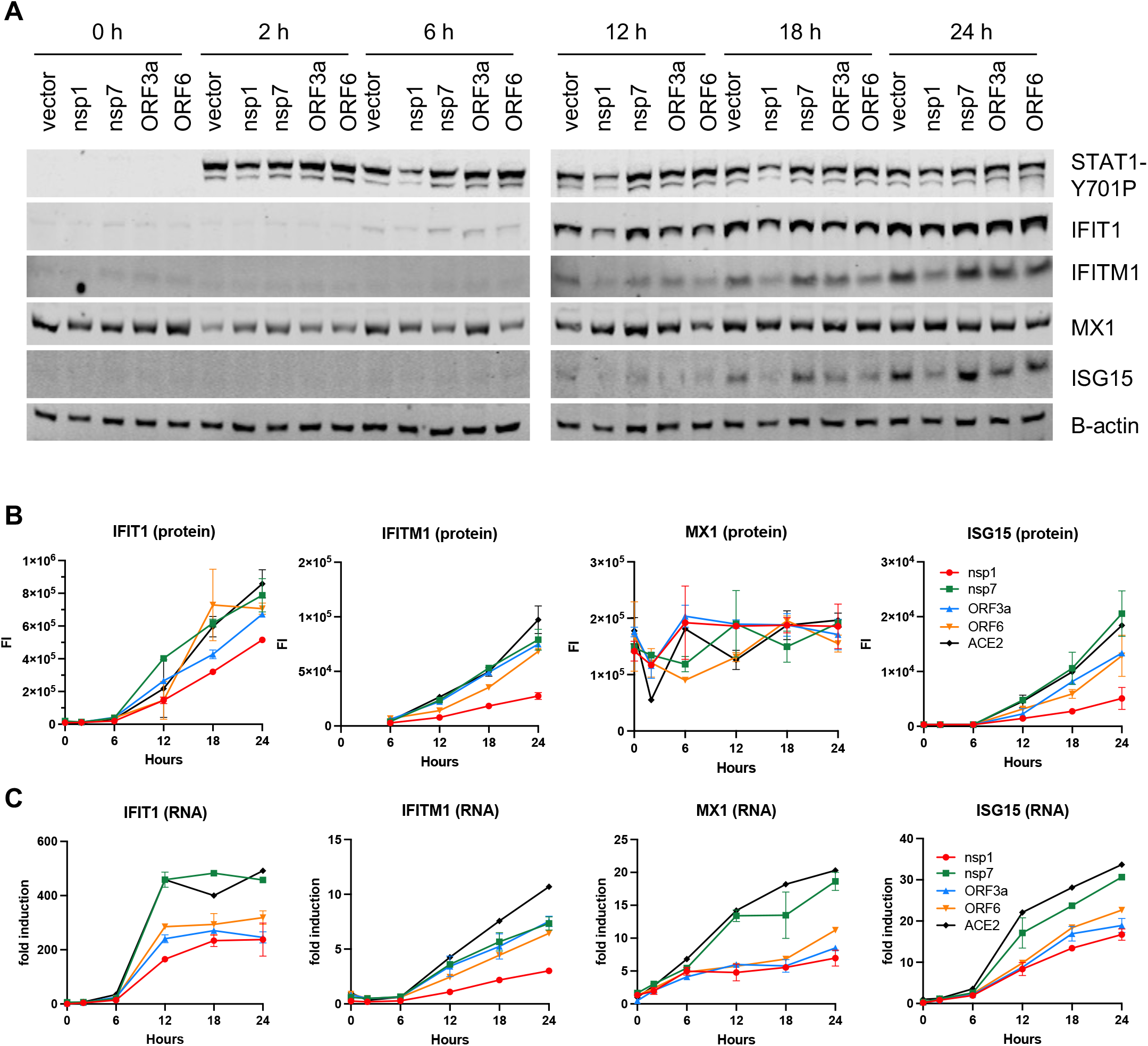
SARS-CoV-2 proteins block the type-I IFN response at different stages. HEK293T cells were transfected with nsp1, nsp7, ORF3a and ORF6 expression plasmids and treated with 1000 u of IFN-alpha. Cells were analyzed for ISG induction by immunoblotting (A, B) and q-RT-PCR (C). Data are derived from two independent experiments. Error bars in B, C show the mean.

## DISCUSSION

Here we utilized ribosome profiling (ribo-seq) coupled with RNA-seq to study the translational events that regulate viral gene expression and host responses over the course of SARS-CoV-2 infection. SARS-CoV-2 replicates rapidly, with viral RNAs constituting the great majority of the total mRNA pool soon after infection. Our data show that viral mRNA abundance is the main determinant of viral protein expression and SARS-CoV-2 mRNAs sequester ribosomes from the translating pool by competition, simply outnumbering the host counterparts. Notwithstanding certain viral mRNAs (i.e. those encoding S, E and ORF1ab) were translated modestly more efficiently than others. While the overall conclusions are similar, another study found that ORF1ab was less efficiently translated compared with other viral mRNAs (Finkel et al., 2020), which we ascribe to possible differences in read depth (with our study having substantially higher read depth within ORF1ab), RNA-seq approaches and infection conditions and cell lines.

We observed that, in contrast to HIV-1, SARS-CoV-2 RNA employs a highly efficient frameshifting strategy to facilitate virus replication. Ribosomal frameshifting requires a heptanucleotide slippery sequence and a RNA pseudoknot, generally a H-type, positioned six to eight nucleotides downstream (Giedroc and Cornish, 2009). Multiple models for ribosomal frameshifting posit that the ribosome pauses at the slippery sequence upon encountering the pseudoknot, which is resistant to unwinding (Farabaugh, 1996, Dinman, 2012). While paused, ribosomes either stay in-frame or slip - 1 nt before resuming translation. A corollary of this notion is that the stimulatory structure, in turn, enhances frameshifting efficiency by promoting ribosomal pausing. In line with this, we observed a local increase in ribosome density overlapping the slippery sequence for both SARS-CoV-2 and HIV-1 (Fig. 1E, **Table S3**), which is also supported by recent structural studies of the ribosome-bound SARS-CoV-2 frameshifting element (Bhatt et al., 2021). Notably, in the case of HIV-1 the increase density of ribosomes extended 100 nucleotides upstream of the frameshifting site, suggesting an alternative frameshifting structure that includes upstream sequences or steric hindrance imposed by the pseudoknot structure. Ribosome density downstream of the frameshifting site within the SARS-CoV-2 ORF1b was high with ribosomes continuing into the ORF1b frame >50% of the time, suggesting the comparably high efficiency of SARS-CoV-2 frameshifting relative to HIV-1 in spite of ribosomal pausing.

We hypothesize that both sequence-specific and structural features contribute to SARS-CoV-2 frameshifting efficiency. It is thought that HIV-1 has a particularly slippery sequence (UUUUUUA) as compared to SARS-CoV-2 (UUUAAAC), which may underlie this difference (Giedroc and Cornish, 2009). In addition, structures downstream of the slippery sequence may have an impact (Plant and Dinman, 2008). For example, the HIV-1 frameshifting element is predicted to have a simpler pseudoknot (Chang et al., 1999, Parkin et al., 1992, Brierley and Dos Ramos, 2006, Huang et al., 2014) or hairpin loop structure (Mouzakis et al., 2013, Staple and Butcher, 2005). In particular, previous studies suggest that frameshifting efficiency positively correlates with the mechanical stability and thermodynamic stability of the pseudoknot and stem loop, respectively (Hansen et al., 2007, Chen et al., 2009, Bidou et al., 1997). In addition, host proteins can also affect frameshifting. Of note, an interferon-stimulated gene (ISG) product, known as C19orf66 (Shiftless), has recently been demonstrated to impair HIV-1 replication through inhibition of HIV-1 programmed frameshifting (Wang et al., 2019). Altogether, our data suggest that SARS-CoV-2 and HIV-1 frameshifting occurs through distinct mechanisms. It remains to be determined how distinct elements within the frameshifting site affect and whether other viral or cellular proteins are involved in modulating the frameshifting efficiencies of these viruses. In addition to frameshifting, we demonstrate that alternative, non-canonical translational start sites internal to several viral genes such as S, E and M, can result in truncated isoforms or short peptides (**Table S4**). Relevance of such findings to SARS-CoV-2 pathogenesis will depend on the development of reverse-genetically modified SARS-CoV-2 strains.

Our study provides an in depth picture of how host cell responses to SARS-CoV-2 are regulated at a post-transcriptional level. For example, in the highly permissive Vero E6 cells, we observed upregulation of proinflammatory chemokines as early as 6 hpi followed by a more delayed induction of ISGs, a finding in line with previous observations in immortalized lung cell lines (Blanco-Melo et al., 2020). However, the increase in transcript abundance did not correlate with higher levels of translation and the great majority of the innate immune response genes appeared to be translated at a low efficiency (Fig. 6A-D, **Table S17**). Apart from this specific effect on, we did not observe a global shutdown of host mRNA translation and most cellular mRNAs were translated proportional to their mRNA abundance.

Translational repression of innate immune genes was less apparent in the complex setting of primary HBECs grown at ALI though several chemokine ligands and IFN-B trended to be less efficiently translated (Fig. 6E-H, **Table S18**). The potential factors that underlie the difference between Vero E6 cells and HBEC-ALI cultures are many-fold. First, Vero E6 cells, as well as other cell line models broadly used in the field (i.e. ACE2 overexpressing cell lines), are unusually permissive to infection allowing quick accumulation of viral proteins with established effects on host mRNA degradation and translation. Second, majority of published models for SARS-CoV-2 infection have utilized cancer-derived cell lines (i.e. Calu-3, A549, Caco-2, Huh7) that often lack kay arms of innate immunity and/or overexpress ACE2, which also enhances rapid SARS-CoV-2 replication. In fact, it is apparent in the HBEC-ALI model that viral translation, and therefore accumulation of viral proteins, may be overall more restricted compared with the highly permissive Vero E6 cells. Third, HBEC-ALI model is composed of other cell types (i.e. basal, club and BC/club cells) that do not express ACE2, and hence are not as efficiently infected by SARS-CoV-2, though there is some evidence that cell tropism can expand to these cells at later stages of SARS-CoV-2 replication (Ravindra et al., 2021). Thus, it is possible that the observed upregulation of inflammatory and innate immune genes takes place in the uninfected bystander cells that do not express viral proteins, a finding consistent with recent scRNA-seq studies (Ravindra et al., 2021). Finally, it is possible that the observed translational repression of innate immune factors may be cell type specific and dependent on high degree of infection. For example, our recent RNA-seq and proteomics studies in the H522 lung adenocarcinoma cells, where SARS-CoV-2 infection progresses slowly, did not reveal a major translational repression of mRNAs encoding host defense factors (Puray-Chavez et al., 2021).

The apparent low translation efficiency of host response mRNAs in Vero E6 cells may be mediated by the SARS-CoV-2 protein NSP1, which associates tightly with the 40S ribosomal subunit as well as non-translating 80S ribosomes to prevent binding of capped mRNA and thus inhibit the formation of the translation initiation complex (Schubert et al., 2020, Banerjee et al., 2020, Thoms et al., 2020, Lapointe et al., 2021), much like its SARS-CoV counterpart (Narayanan et al., 2015). In addition, there is increasing evidence that ectopic expression of NSP1 can alter host mRNA translation (Rao et al., 2021). Given the high abundance of ribosomes in the cell, whether physiologically relevant concentrations of NSP1 is sufficient to induce a global block in mRNA translation remains unknown. For example, even in cells overexpressing NSP1, we did not observe a major translational block to ISG induction (Fig. 7). Rather, NSP1 expression blocked STAT1 phosphorylation and subsequently reduced transcriptional induction of ISGs. Thus the observed translational repression of ISGs in the heavily-infected Vero E6 cells is likely due to a combination of viral mRNAs dominating the cellular mRNA pool, other viral proteins such as NSP1 and possibly due to reduced translation initiation due to cellular stress induced by SARS-CoV-2. Finally, we cannot rule out the possibility that the translational suppression of innate immune genes is also contributed by the host’s attempt to curb viral replication, including members of the IFIT family with known functions in translation inhibition (Hyde and Diamond, 2015, Fensterl and Sen, 2015, Reynaud et al., 2015, Daffis et al., 2010, Diamond and Farzan, 2013). Future studies are warranted to empirically test these possibilities and define the mechanism of apparent innate immune suppression.

While COVID-19 pathogenesis is in part due to virus-induced destruction of infected cells, elevated production of inflammatory mediators and the virus-induced immunopathology are thought to play a big role in SARS-CoV-2-induced lung injury (Channappanavar and Perlman, 2017, Perlman and Dandekar, 2005). Our findings suggest that immune responses in actively infected cells may be dampened or delayed for SARS-CoV-2 to efficiently replicate and release viral progeny. As such, it is possible that the elevated levels of inflammatory mediators *in vivo* is due to by-stander cells or infection of immune cell subsets, such as monocytes and macrophages, that are less permissive to SARS-CoV-2 but can sense and respond to infection by secretion of immune modulatory molecules (Jafarzadeh et al., 2020).

Taken together, we provide novel insight into and a rich resource on how translational regulation shapes SARS-CoV-2 replication and host responses. Our finding that induction of inflammatory and innate immune responses can be limited at the level of mRNA translation provides a paradigm shifting mechanism of how SARS-CoV-2 can encounter immune responses. Modulation of viral RNA structures and proteins that regulate mRNA translation will undoubtedly provide a unique avenue for therapeutic development. Together, our study provides an in-depth picture of translationally regulated events in SARS-CoV-2 replication and reveal that impairment of host mRNA translation may allow SARS-CoV-2 to evade host immunity.

## ACKNOWLEDGEMENTS

We would like to thank members of the Diamond lab for reagents and support. This work was supported by Washington University startup funds for SBK, Andrew and Virginia Craig faculty fellowship for SBK, R21 AI145669 to SBK and JD, National Science Foundation Graduate Research Fellowship under Grant No. DGE-1745038 to KV, Stephen I. Morse postdoctoral fellowship to MPC, Helen Hay Whitney Foundation postdoctoral fellowship to J.B.C., and the Dorothy R. and Hubert C. Moog Professorship to SLB. All NGS data (48 sets of ribo-seq and 48 sets of matching RNA-seq) are deposited in the GEO database under GSE158930.

## AUTHOR CONTRIBUTIONS

MPC, HRV, AH, TH, SGP, JBC, SBK conducted the experiments. KT, NL, YW, YL, WY conducted the bioinformatics analysis. HRV and SBK wrote the original manuscript with input from MSD, SLB, JD, JBC.

## MATERIALS AND METHODS

### Chemicals and reagents

Standard laboratory chemicals were obtained from reputable suppliers such as Sigma-Aldrich. Cycloheximide (CHX) was obtained from Sigma, dissolved in ethanol and stored at −20°C. Harringtonine (HT) was purchased from LKT Laboratories, Inc., resuspended in DMSO and stored in aliquots of 2 mg/mL at −20°C.

### Plasmids and viruses

A proviral plasmid encoding the full length HIV-1 genome was obtained from NIH AIDS Reagents. HIV-1 stocks were generated by transfection of Human embryonic kidney cell line, HEK293T, with proviral plasmids and collection of cell culture supernatants two days after. Viruses were treated by DNase to avoid plasmid carryover and concentrated by Lenti-X concentrator. HIV-1 stocks were titered on TZM-bl cells by conventional methods. CD4^+^ T-cells activated for 4-5 days were used for HIV-1 infections. SARS-CoV-2 strain 2019-nCoV/USA-WA1/2020 was obtained from Natalie Thornburg at the Centers for Disease Control and Prevention (CDC), propagated in Vero CCL-81 cells and titrated on Vero E6 cells by plaque-forming assays. SARS-CoV-2 Neon-green reporter virus has been explained before (Xie et al., 2020) and was propagated and titered similarly. Mammalian expression plasmids encoding SARS-CoV-1 genes (NSP1, NSP7, ORF3a, ORF6) were obtained from BEI resources and propagated as recommended.

### Cells and infections

HEK293T and TZM-bl cells were obtained from ATCC and NIH AIDS Reagent Program respectively and were maintained in Dulbecco’s Modified Eagle’s Medium (DMEM) (high glucose), supplemented with 10% fetal bovine serum (FBS) in a humidified incubator at 37°C with 5% CO_2_. For isolation of primary CD4^+^ T-cells, buffy coats (from anonymous healthy blood donors from Mississippi Blood Center) were separated by Ficoll and CD4^+^ T-cells purified using RosetteSep Human CD4+ T-cell enrichment kit (STEMCELL Technologies). CD4^+^ T-cells cells were maintained in Roswell Park Memorial Institute 1640 medium (RPMI) supplemented with 10% heat-inactivated FBS and L-glutamine. Activation of CD4^+^ T cells was achieved by addition of 25 U/ml of interleukin-2 (IL-2) and CD4^+^ T-cell activation Dynabeads (Life Technologies). CD4^+^ T-cells activated for 4-5 days were used for HIV-1 infections. Vero CCL-81 and Vero E6 were cultured in DMEM supplemented with 10% FBS and 10 mM HEPES pH 7.4. For SARS-CoV-2 infections, Vero E6 cells were incubated with SARS-CoV-2 inoculum in DMEM-supplemented with 2% FBS for an hour in a humidified incubator at 37°C, after which the initial inoculum was removed and replaced by cell culture media.

Primary human bronchial epithelial cells (HBECs) grown at air-liquid interface (ALI). Human airway epithelial cells were isolated from surgical excess of tracheobronchial segments of lungs donated for transplantation as previously described and were exempt from regulation by US Department of Health and Human Services regulation 45 Code of Federal Regulations Part 46 (Horani et al., 2012). Tracheobronchial cells were expanded in culture, seeded on supported membranes (Transwell; Corning, Inc.), and differentiated using ALI conditions as detailed before (You et al., 2002, Horani et al., 2018).

### Immunofluorescence

Infected Vero E6 cells and HBECs were fixed with 4% paraformaldehyde for 20 min at room temperature, followed by permeabilization using 0.5% Tween-20 in PBS for 10 min. Cells were blocked with 1% bovine serum albumin (BSA) and 10% FBS in 0.1% Tween-20 PBS (PBST) for 1 h prior to staining with a rabbit polyclonal anti SARS-CoV-2 nucleocapsid antibody (Sino Biological Inc. catalog # 40588-T62) diluted 1:500 and incubated overnight at 4°C. The following day, cells were stained with an Alexa Fluor 488-conjugated goat anti-rabbit secondary antibody (Invitrogen) at 1:1000 dilution, counter-stained with DAPI and imaged by immunofluorescence microscopy.

### RNA in situ hybridization

Primary-culture human airway epithelial cells were fully differentiated at air-liquid interface on supported plastic membranes (Transwell, Corning). Cells were fixed by immersion of the Transwell membrane in methanol:acetone (50:50% volume) at −20 °C for 20 min followed by 4% paraformaldehyde at room temperature for 15 min. Cells were washed three times with phosphate buffered saline (PBS) and stored at 4 °C. Prior to probing for vRNA, membranes containing cells were cut from plastic supports, divided into 4 pieces, and placed in a fresh 48-well plate. RNA detection was performed using the manufacturer protocol for RNAscope fluorescent in situ hybridization (RNAscope Multiplex Fluorescent v2 Assay kit, Advanced Cell Diagnostics). Briefly, cells on membranes were treated with 3% hydrogen peroxide for 10 min at room temperature, washed with distilled water, then treated with protease III solution, diluted 1:15 in PBS, for 10 min in a humidified hybridization oven at 40 °C. The cells were then washed with PBS and incubated for 2 hr at 40 °C with manufacturer designed anti-sense probes specific for SARS-CoV-2 positive strand S gene encoding the spike protein (RNAscope Probe-V-nCoV2019-S, cat# 848561) or ORF1ab (RNAscope Probe-V-nCoV2019-orf1ab-O2-sense-C2 cat # 854851-C2). The probes were visualized according to the manufacturers’ instructions by incubation with RNAscope amplifiers, horseradish peroxidase, and fluorescent label (Opal fluorophores, Perkin-Elmer). Membranes were mounted on glass slides using anti-fade medium containing DAPI (Fluoroshield, Sigma-Aldrich). Images were obtained using a 5000B Leica microscope equipped with a charge-coupled device camera (Retiga 200R) interfaced with QCapture Pro software (Q Imaging).

### Ribosome profiling

Ribosome profiling (Ribo-seq) was performed as described before with the following modifications (Ingolia et al., 2009, Ingolia et al., 2012). Mock- and HIV-1- or SARS-CoV-2-infected cells were treated with complete cell culture media supplemented with 0.1 mg/mL CHX for 1 min at room temperature followed by one round of wash in ice-cold PBS supplemented with 0.1 mg/mL CHX. Cells were lysed in 1X mammalian polysome lysis buffer (20 mM Tris·HCl (pH 7.4), 150 mM NaCl, 5 mM MgCl_2_, 1% Triton X-100, 0.1% NP-40, 1 mM DTT, 10 units of DNase I, with 0.1 mg/mL CHX). The cells were then triturated by repeated pipetting and incubated with lysis buffer for at least 20 min to ensure virus inactivation. Lysates were centrifuged for 10 min at ≥20,000 g at 4°C for clarification. The supernatants were split into multiple aliquots, with SDS added to one aliquot to a final concentration of 1% for downstream RNA-seq sample preparation, and flash frozen in a 70% ethanol/dry ice bath or directly placed at −80°C. RNA extracted from lysates were subjected to Bioanalyzer RNA-Nano analysis. RNA integrity number (RIN) of 8 and above (max RIN = 10) is considered “intact RNA”. Lysates were treated with RNase I (5U/OD_260_) and ribosome-protected fragments were isolated via centrifugation through Microspin S-400 HR columns (GE Healthcare) and purified using the RNA Clean and Concentrator kit (Zymo Research). Recovered ribosome-bound fragments (RBFs) are then subjected to rRNA depletion using RiboZero beads from the TruSeq Stranded Total RNA Library Prep Gold kit (Illumina) and purified using Zymo RNA Clean and Concentrator kit. Fragments were then end-labeled with γ-^32^P-ATP using T4 polynucleotide kinase (NEB), separated on 15% TBE-Urea gels and visualized by autoradiography. RNA fragments of ∼30 nt were excised from the gels and purified as detailed before in 400 μL of 0.4 M NaCl supplemented with 4 μL SUPERaseIN. 3’ and 5’ adapters were sequentially ligated as in a previously described protocol (Kutluay et al., 2014, Kutluay and Bieniasz, 2016), reverse transcribed and PCR amplified. We acknowledge that our ligation-based library generation protocol may introduce biases towards inserts containing distinct nucleotides at the 5’ and 3’ end. Indeed, we found a modest preference towards Us and Cs in the first position and Gs and Cs in the last position of inserts. Libraries were then sequenced on HiSeq-2000 or NextSeq 500 platforms (Illumina) at the Genome Technology Access Center or the Edison Family Center for Genome Sciences & Systems Biology, respectively, at Washington University School of Medicine. All ribo-seq and RNA-seq data were deposited on GEO database under GSE158930.

### RNA-seq

An aliquot of cell lysates harvested from ribosome profiling (Ribo-seq) experiments above was processed in parallel for RNA-seq using TruSeq Stranded mRNA library prep (Illumina) following extraction using Zymo RNA-Clean and Concentrator (5) kit. RNA-seq libraries were constructed using TruSeq RNA single-index adapters and deep sequenced as above at Washington University in St. Louis.

### Data analysis

All of the data analysis pipelines used in this study are deposited to GitHub under kutluaylab. Below we detail the salient steps of data analyses:

#### Mapping

Generated RNA-seq and Ribo-seq data were analyzed by publicly available software and custom scripts. In brief, for Ribo-seq, reads were separated based on barcodes and the adapters trimmed using BBDuk (http://jgi.doe.gov/data-and-tools/bb-tools/) and FastX Toolkit (http://hannonlab.cshl.edu/fastx_toolkit/). The resulting reads were mapped to the viral genome/transcriptome using the Bowtie aligner (Langmead et al., 2009) (mapping criteria -v 1, -m 10), and to the African green monkey (AGM) (*Chlorocebus sabaeus*) or human genomes (hg19) in STAR (Dobin et al., 2013) (mapping criteria FilterMismatchNoverLmax 0.04). For ribo-seq reads that map to the SARS-CoV-2 genome, reads were additionally collapsed to minimize PCR overamplification artifacts. For AGM/human alignments, reads were first mapped to rRNA to remove any rRNA-derived reads not completely removed by depletion kits and to the SARS-CoV-2 genome to remove virally derived reads. The remaining reads were then first mapped to the SARS-CoV-2 genome (to remove viral reads) and then to the AGM/human genomes. RNA-seq reads were similarly mapped using STAR, although the rRNA alignment step was omitted. For AGM/human alignments, mapped reads were annotated using the featureCounts package and GTF files freely available from NCBI and Ensembl.

#### Statistical Analysis

Differential gene expression analysis was carried out using the edgeR package (Robinson et al., 2010). Considering that virally derived sequences quickly dominated the host mRNA pool, for differential gene expression of host mRNAs, library sizes were normalized relative to reads that map only to host mRNAs. Efforts in this area focused on determining upregulated genes using individual Ribo-seq and RNA-seq experiments, as well as the analysis of log_2_-fold change differences between Ribo-seq and RNA-seq to discover translationally regulated genes. These same files and packages were also used to generate quality control plots and graphics highlighting differentially expressed genes. The calculation of translational efficiency involved normalizing counts to account for library size in edgeR to generate log_2_ counts-per-million (log_2_CPM) estimates for each gene in Ribo-seq and RNA-seq, and subtracting log_2_CPM RNA-seq from log_2_CPM Ribo-seq to provide an estimate of the difference in expression level between Ribo-seq and RNA-seq for a given gene.

Downstream analysis of sets of differential genes involved the use of goseq (Young et al., 2010) and KEGGREST R packages (Tenenbaum, 2020). Annotations of 5’UTRs, CDSs and 3’UTRs were retrieved and repetitive low-complexity elements were removed. The R package riboWaltz (Lauria et al., 2018) and the Ribo-TISH package (Zhang et al., 2017) were utilized to determine the location of ribosomal P-sites with respect to the 5’ and 3’ end of reads, as well as illustrating triplet periodicity and determining the percentage of reads within each frame in CDS and UTR. Finally, the metagene R package (Beauparlant, 2020) was applied to generate an aggregate analysis of ribosomal density downstream of start codons and upstream of stop codons in the corresponding genome.

Alternative TIS sites in both host and viral reads were found using the Ribo-TISH package (Zhang et al., 2017). For viral TIS, analysis was carried out in the ‘predict’ mode comparing samples mock-treated or treated with harringtonine at each timepoint (with replicates). This was replicated for host analysis, although with the additional step of analysis in the ‘diff’ mode to predict TIS differentially regulated between infected and uninfected cells.

#### Cluster analysis

RNA-seq and ribo-seq logCPM values were each converted to per-gene z-scores. Consensus clustering was then performed with the R ConsensusClusterPlus package (Wilkerson and Hayes, 2010). The non-defaults settings used were: reps=50, innerLinkage=”complete”, and finalLinkage=”ward.D2”. The optimal number of clusters was chosen by manual inspection of clustering quality for consensus matrices with k=1-12.

#### Gene set enrichment analysis

Over-representation of biological gene sets in individual temporal gene clusters for RNA-seq and ribo-seq data were investigated using the R clusterProfiler package and enricher function (Yu et al., 2012). Gene sets were downloaded from the MSIG data bank via the msigdbr R-project package, including “Hallmark” and “GO:BP” (Liberzon et al., 2011, Liberzon et al., 2015, Jassal et al., 2020). Gene sets were considered significantly enriched in a cluster if adjusted q-values were < 0.05.

#### Viral Counts

Viral read density plots were generated using the SAM file from viral genome alignment. The SAMtools (Li et al., 2009) package was used to create an mpileup file containing information about the read density, strandedness, mapping quality, and nucleotide identity at each position. Custom scripts (deposited at GitHub under kutluaylab) then were utilized to create files providing only the nucleotide identity and number of counts at each position for both sense and antisense reads. These were then visualized by scripts written in R.

As SARS-CoV-2 generates chimeric subgenomic mRNAs (sgRNAs) in addition to its genomic RNA (gRNA), featureCounts could not be used to accurately estimate viral gene counts from RNASeq due to the presence of nested 3’ sequences. Therefore, in order to visualize and enumerate such chimeric sequences the BWA aligner (Li and Durbin, 2009) was used in ‘mem’ mode on viral RNASeq reads. After generating this alignment using the default parameters and same reference SARS-CoV-2 FASTA file as above, chimeric reads were isolated by searching for all reads containing the ‘SA’ tag and the SARS TRS sequence, ‘AAACGAAC’. SARS-CoV-2 gRNAs were extracted by searching for all reads containing the first 15-20 bases of the ORF1A coding sequence (CDS), as these sequences would only be present in full-length SARS-CoV-2 genomes. This provided the sequences and alignment locations of the chimeric and genomic reads, which were then visualized using R. For sgRNAs, the viral gene corresponding to each transcript was determined by locating the CDS with the nearest downstream start site. This data, together with the number of gRNAs was used to calculate relative percentages of viral transcripts and, together with the total number of mapped viral reads, allowed for the tabulation of viral gene counts at each time point. For ribosome profiling data, featureCounts was used to enumerate the number of viral reads, as ribosomes only translate the first gene on each transcript and so footprints from nested 3’ gene were low enough to be negligible.

## SUPPLEMENTARY FIGURE LEGENDS

**Figure S1. SARS-CoV-2 infection of Vero E6 cells (Supplemental to** Figure 1). Vero E6 cells were infected at an MOI of 2 pfu/cell. (A) Infected cells were processed at 2, 6, 12 and 24 hpi for immunofluorescence microscopy using an antibody against the viral N protein. Scale bar = 100 uM. (B) The integrity of the RNA samples used in RNA-seq/ribo-seq experiments was analyzed by Bioanalyzer RNA-nano. RNA integrity number (RIN) is indicated on the plots. RIN values range from 0-10, with RIN=10 indicative of intact RNA in samples.

**Figure S2. Quality control of Ribo-seq libraries derived from SARS-CoV-2-infected Vero E6 cells (Supplemental to** Figure 1). Vero E6 cells infected as in Fig. 1A were processed for ribo-seq as detailed in Materials and Methods. (A) Length distribution of ribo-seq-derived reads mapping to the African green monkey (AGM) and SARS-CoV-2 transcriptomes are shown for three independent replicate libraries. (B) Number or reads mapping to 5’UTR, CDS and 3’ UTRs of annotated AGM genes in matching RNA-seq and ribo-seq libraries are shown.

**Figure S3. P-site analysis of reads that map to cellular transcripts derived from ribo-seq experiments done on SARS-CoV-2-infected Vero E6 cells (Supplemental to** Figure 1). Ribo-seq libraries were derived from SARS-CoV-2-infected Vero E6 cells (as in Fig. 1A) were analyzed as detailed in Materials and Methods. (A) Meta-profiles showing the periodicity of ribosomes along the AGM transcripts at the genome-wide scale from independent replicate samples. (B) Enrichment of P-sites in different frames for reads of varying length mapping to 5’ UTR, CDS and 3’ UTRs are shown.

**Figure S4. P-site analysis of reads that map to viral transcripts derived from ribo-seq experiments done on SARS-CoV-2-infected Vero E6 cells (Supplemental to** Figure 1). Ribo-seq-derived reads obtained from SARS-CoV-2-infected (MOI: 2 i.u./cell, 6hpi) Vero E6 cells were mapped to the viral transcriptome and analyzed as detailed in Materials and Methods. Different quality profiles/metrics for RPFs uniquely mapped to annotated protein-coding regions for the three replicate data sets are shown. The data corresponding to the first, second and third reading frame are colored in red, green blue, respectively. Each row shows the RPFs with indicated length. Column 1: distribution of RPF 5′ end across three reading frames in all annotated codons; showing the fraction of RPF counts from dominant reading frame. Column 2: distribution of RPF 5′ end count near annotated TISs; showing estimated P-site offset and the ratio between the RPF counts at the annotated TISs and the sum of the RPF counts near the annotated TISs (from −1 to +1 relative to the annotated TISs) after P-site offset correction. Column 3: distribution of RPF 5′ end count near annotated stop codon. Column 4: RPF count profile throughout protein-coding regions across three reading frames.

**Figure S5. Ribo-seq in SARS-CoV-2-infected cells (Supplemental to** Figure 1). Vero E6 cells infected with SARS-CoV-2 at an MOI of 2 pfu/cell were processed for RNA-seq and ribo-seq as detailed in Materials and Methods. (A) RNA-seq (log-scale) and (B) Ribo-seq (linear scale) counts along the viral genome across various time points. Schematic diagram of SARS2 genome features shown above is co-linear. (Also see **Table S3**)

**Figure S6. Ribosome occupancy on SARS-CoV-2 transcripts during high MOI infection (Supplemental to** Figure 1). (A) Ribo-seq and RNA-seq data derived from experiments described in Figure 1 were plotted to demonstrate the number of reads that map to SARS-CoV-2 transcripts at 2 hpi (Also see **Table S3**). Length distribution of reads that map to the viral genome is shown on the right for each replicate. (B) Predicted secondary structure of the SARS-CoV-2 frameshifting element is shown.

**Figure S7. Frame information of alternative translation initiation sites derived from ribo-seq experiments done in the presence of harringtonine.** Vero E6 cells were infected with SARS-CoV-2, MOI: 2 pfu/cell as in Fig 1A. Frames of each read that map to the SARS-CoV-2 genome is shown following P-site analysis. Alternative translation initiation sites are indicated on the plot (Also see **Table S4**).

**Figure S8. Quality control of Ribo-seq libraries derived from primary CD4+ T-cells infected with HIV-1 (Supplemental to** Figure 2). CD4+ T-cells isolated from two independent donors infected HIV-1 were processed for ribo-seq as detailed in Materials and Methods. (A) Length distribution of ribo-seq-derived reads mapping to the human transcriptome and HIV-1 genome are shown for independent replicates. (B) Enrichment of P-sites in different frames for reads of varying length mapping to 5’ UTR, CDS and 3’ UTRs are shown. (C) Meta-profiles showing the periodicity of ribosomes along the human transcripts at the genome-wide scale from independent replicate samples. (D) Number or reads mapping to 5’UTR, CDS and 3’ UTRs of annotated human genes in matching RNA-seq and ribo-seq libraries are shown.

**Figure S9. SARS-CoV-2 infection of primary HBECs grown at ALI (supplemental to** Figure 3). (A) Primary HBEC cultures grown at ALI were infected with SARS-CoV-2-mNG at an MOI of 1 pfu/cell and imaged live by epifluorescence microscopy at 48, 72 and 96hpi. (B) Primary HBEC cultures were infected with SARS-CoV-2 at an MOI of 1 pfu/cell and fixed at 96 hpi. Cells were probed with RNAScope probes against sense- and anti-sense SARS-CoV-2 transcripts, and imaged by immunofluorescence microscopy with a 4X objective.

**Figure S10. RNA-seq in SARS-CoV-2-infected cells (Supplemental to** Figure 3). Primary HBEC cultures grown at ALI were infected with SARS-CoV-2 at an MOI of 1 pfu/cell and processed for RNA-seq as detailed in Materials and Methods. RNA-seq (log-scale) counts along the viral genome across various time points. Schematic diagram of SARS2 genome features shown above is co-linear. (Also see **Table S10**).

**Figure S11. Quality of ribo-seq experiments from SARS-CoV-2-infected primary HBECs grown at ALI (supplemental to** Figure 3). Primary HBEC cultures were infected with SARS-CoV-2 at an MOI of 1 pfu/cell as in Figure 3 and processed for ribo-seq at the indicated times post-infection. (A) The integrity of the RNA samples used in RNA-seq/ribo-seq experiments was analyzed by Bioanalyzer RNA-nano. RNA integrity number (RIN) is indicated on the plots. RIN values range from 0-10, with RIN=10 indicative of intact RNA in samples. (B) Length distribution of ribo-seq-derived reads mapping to the human and SARS-CoV-transcriptome are shown.

**Figure S12. Quality of ribo-seq experiments from SARS-CoV-2-infected primary HBECs grown at ALI (supplemental to** Figure 3). Primary HBEC cultures were infected with SARS-CoV-2 at an MOI of 1 pfu/cell as in Figure 3 and processed for ribo-seq at the indicated times post-infection. Enrichment of P-sites in different frames for reads of varying length mapping to 5’ UTR, CDS and 3’ UTRs are shown.

**Figure S13. Metaprofiles and P-site analyses of ribo-seq reads derived from SARS-CoV-2-infected primary HBEC cells (Supplemental to** Figure 3). (A) Meta-profiles showing the periodicity of ribosomes along the human transcripts at the genome-wide scale from independent ribo-seq libraries derived from primary HBECs at the indicated times post-infection. (B) Different quality profiles/metrics for RPFs uniquely mapped to annotated protein-coding regions for cellular and viral transcripts from representative data sets are shown. Because the number of reads that map to viral transcripts were relatively low at a given time point post-infection, virally mapping reads across all time points are shown from one of the four representative experiments. The data corresponding to the first, second and third reading frame are colored in red, green blue, respectively. Each row shows the RPFs with indicated length. Column 1: distribution of RPF 5′ end across three reading frames in all annotated codons; showing the fraction of RPF counts from dominant reading frame. Column 2: distribution of RPF 5′ end count near annotated TISs; showing estimated P-site offset and the ratio between the RPF counts at the annotated TISs and the sum of the RPF counts near the annotated TISs (from −1 to +1 relative to the annotated TISs) after P-site offset correction. Column 3: distribution of RPF 5′ end count near annotated stop codon. Column 4: RPF count profile throughout protein-coding regions across three reading frames.

**Figure S14. Reproducibility of RNA-seq and ribo-seq experiments in SARS-CoV-2-infected Vero E6 cells (Supplemental to** Figure 4). Vero E6 cells were infected as in Figure 1A and processed for RNA-seq and ribo-seq. (A) Principle component analysis (PCA) of highly expressed genes in RNA-seq and ribo-seq experiments across all time points. (B, C) Biological coefficient of variation (BCV) plots demonstrate the gene level biological variation in RNA-seq (B) and ribo-seq (C) data sets under the indicated conditions.

**Figure S15. Cluster profiles of differentially expressed genes derived from infected Vero E6 cells (Supplemental to** Figure 4). Vero E6 cells were infected as in Figure 4 and differentially expressed genes were clustered based on z-score. Cluster profiles show quantification of each gene across the clusters identified in Figure 4 for RNA-seq (A) and ribo-seq (B) data sets. The colored lines represent quantification of an individual protein whereas the solid black and dashed black lines represent the mean of infected and mock samples, respectively.

**Figure S16. Time-course analysis of differentially expressed genes in response to SARS-CoV-2 infection in Vero E6 cells (Supplemental to** Figure 4). Vero E6 cells were infected as in Figure 1A and processed for RNA-seq and ribo-seq. Differentially expressed genes across different time points from Figure 4 were plotted to individually to demonstrate the time-course progression of differential gene expression in RNA-seq (A) and ribo-seq (B) data sets. Data show the log_2_(cpm) values of genes that are up- or down-regulated greater than 2-fold with FDR<0.05. See also **Table S11, S12**.

**Figure S17. Time-course analysis of differentially expressed genes in response to low MOI SARS-CoV-2 infection in Vero E6 cells (Supplemental to** Figure 4). Vero E6 cells were infected with SARS-CoV-2 at an MOI of 0.1 i.u/cell and differentially expressed genes derived from replicate RNA-seq (A) and ribo-seq (B) libraries were are shown. See also **Table S14.**

**Figure S18. Reproducibility of RNA-seq and ribo-seq experiments in SARS-CoV-2-infected HBECs (Supplemental to** Figure 5). Primary HBEC cells grown at ALI as were infected as in Figure 3A and processed for RNA-seq and ribo-seq. (A) Principle component analysis (PCA) of highly expressed genes in RNA-seq and ribo-seq experiments across all time points. Replicate 1 and 2 are from samples obtained from the first donor, and replicates 3 and 4 are from the second donor. (B, C) Biological coefficient of variation (BCV) plots demonstrate the biological variation in RNA-seq (B) and ribo-seq (C) data sets under the indicated conditions.

**Figure S19. Cluster profiles of differentially expressed genes derived from infected HBEC cells grown at ALI (Supplemental to** Figure 5). Primary HBECs grown at ALI were infected as in Figure 3A and differentially expressed genes were clustered based on z-score. Cluster profiles show quantification of each gene across the clusters identified in Figure 4 for RNA-seq (A) and ribo-seq (B) data sets. The colored lines represent quantification of an individual protein whereas the solid black and dashed black lines represent the mean of infected and mock samples, respectively.

**Figure S20. Time-course analysis of differentially expressed genes in response to SARS-CoV-2 infection in HBECs (Supplemental to** Figure 5). Primary HBEC cells grown at ALI as were infected as in Figure 3A and processed for RNA-seq and ribo-seq. Differentially expressed genes across different time points from Figure 4 were plotted to individually to demonstrate the time-course progression of differential gene expression in RNA-seq (A) and ribo-seq (B) data sets. Data show the log_2_(cpm) values of genes that are up- or down-regulated greater than 2-fold with FDR<0.05. See also **Table S15, S16**.

## SUPPLEMENTARY TABLES

**Table S1.** Ribosome profiling sequencing data statistics from Vero cells infected with SARS-CoV-2 at MOI: 2 pfu/cell

**Table S2.** RNA-seq data statistics from Vero cells infected with SARS-CoV-2 at MOI: 2 pfu/cell.

**Table S3.** Summary of reads mapping to SARS-CoV-2 genome from RNA-seq and ribo-seq derived from infected Vero E6 cells.

**Table S4.** Alternative translation initiation sites in SARS-CoV-2

**Table S5.** Ribosome profiling sequencing data statistics derived from primary CD4+ T-cells infected with HIV-1.

**Table S6.** HIV-1 RNA-seq data statistics derived from primary CD4+ T-cells infected with HIV-1.

**Table S7.** Summary of reads mapping to HIV-1 genome from RNA-seq and ribo-seq derived from infected primary CD4+ T-cells.

**Table S8.** Ribosome profiling sequencing data statistics from SARS-CoV-2 infected HBEC cells grown at ALI.

**Table S9.** RNA sequencing data statistics from SARS-CoV-2 infected HBEC cells grown at ALI.

**Table S10.** Summary of reads mapping to SARS-CoV-2 genome from RNA-seq and ribo-seq derived from infected primary HBEC-ALI cultures.

**Table S11.** Differentially expressed genes and pathways derived from RNA-seq experiments of SARS-CoV-2-infected (high MOI) Vero E6 cells.

**Table S12.** Differentially expressed genes and pathways derived from ribo-seq experiments of SARS-CoV-2-infected (high MOI) Vero E6 cells.

**Table S13.** RNA-seq data statistics from Vero cells infected with SARS-CoV-2 at MOI:0.1 pfu/cell.

**Table S14.** Differentially expressed genes and pathways derived from RNA-seq and ribo-seq experiments of SARS-CoV-2-infected (low MOI) Vero E6 cells.

**Table S15.** Differentially expressed genes and pathways derived from RNA-seq experiments of SARS-CoV-2-infected primary HBECs.

**Table S16.** Differentially expressed genes and pathways derived from ribo-seq experiments of SARS-CoV-2-infected primary HBECs.

**Table S17.** Translation efficiency of DEGs in SARS-CoV-2-infected (high MOI) Vero E6 cells.

**Table S18.** Translation efficiency of DEGs in SARS-CoV-2-infected primary HBECs.

## Notes

### Competing Interest Statement

The authors have declared no competing interest.

### Summary of Updates

More extensive data analysis and new data.

