## Supplementary Figs for "The translational landscape of SARS-CoV-2 and infected cells"

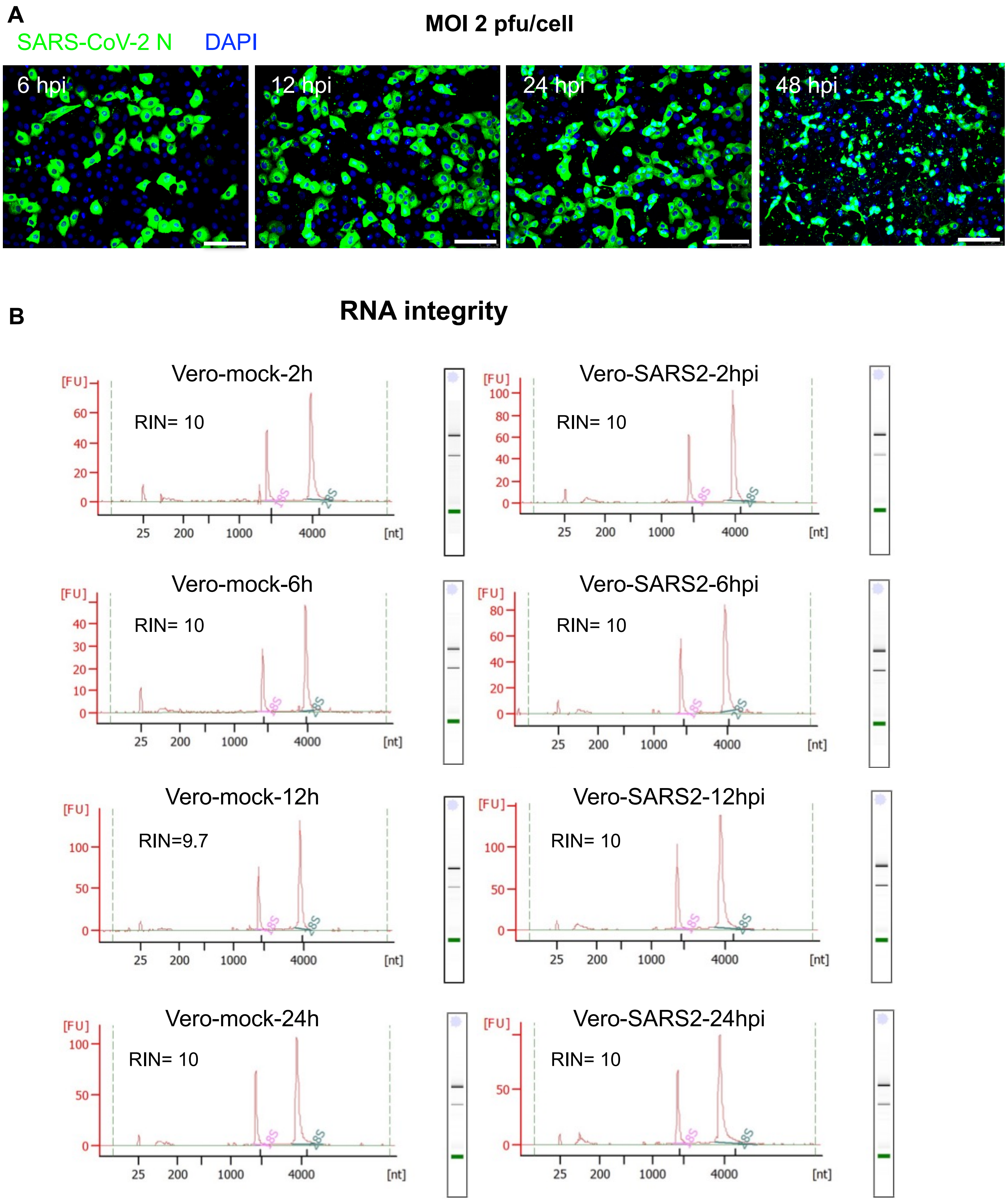
A
SARS-CoV-2 N
DAPI
MOI 2 pfu/cell
6 hpi
12 hpi
24 hpi
48 hpi
B
RNA integrity
Vero-mock-2h
RIN= 10
Vero-SARS2-2hpi
RIN= 10
Vero-mock-6h
RIN= 10
Vero-SARS2-6hpi
RIN= 10
Vero-mock-12h
RIN=9.7
Vero-SARS2-12hpi
RIN= 10
Vero-mock-24h
RIN= 10
Vero-SARS2-24hpi
RIN= 10
[FU]
[nt]
25
200
1000
4000
18S
28S

Figure S1

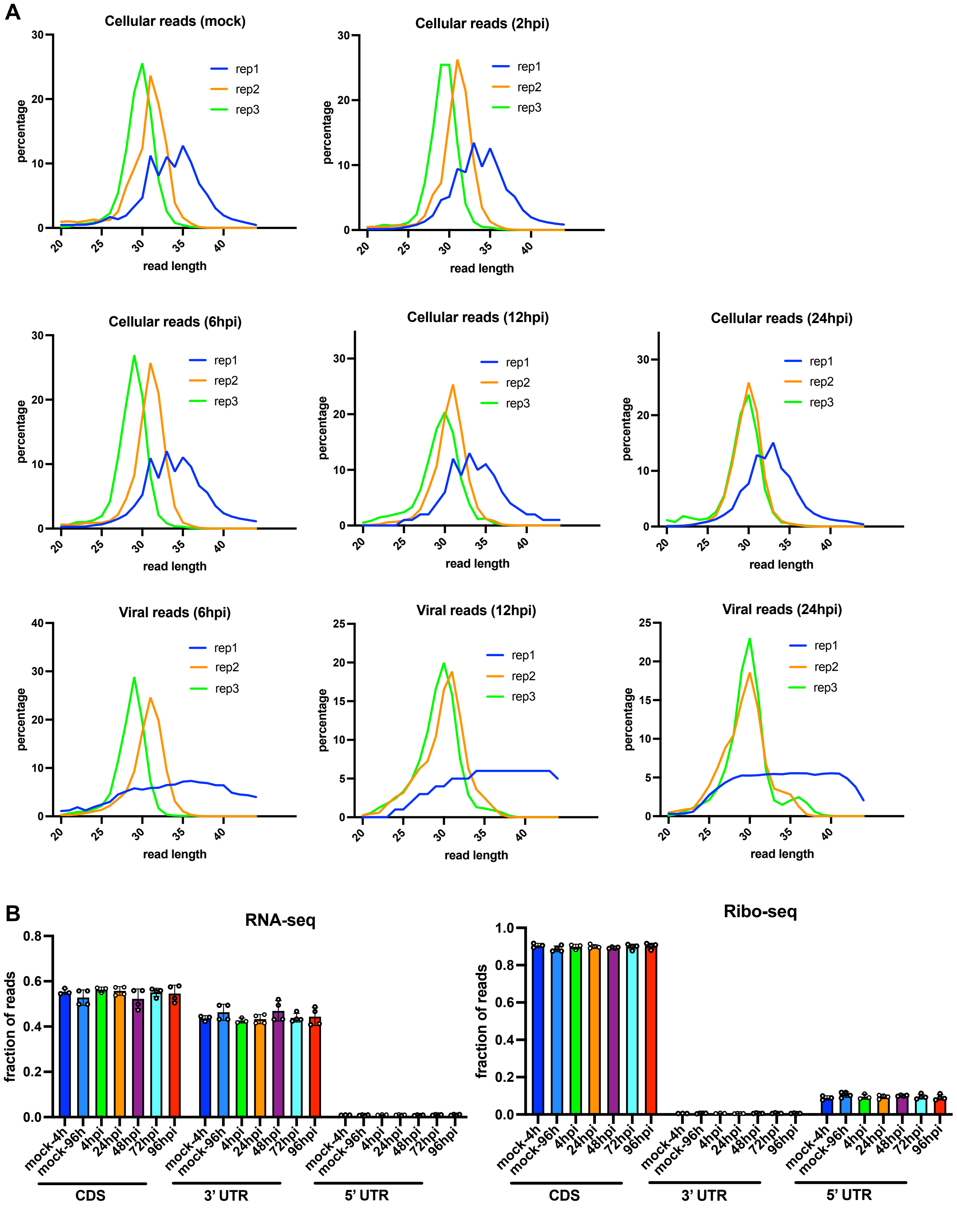

A
Cellular reads (mock)
Cellular reads (2hpi)
Cellular reads (6hpi)
Cellular reads (12hpi)
Cellular reads (24hpi)
Viral reads (6hpi)
Viral reads (12hpi)
Viral reads (24hpi)
rep1
rep2
rep3
percentage
read length
B
RNA-seq
Ribo-seq
fraction of reads
mock-4h
mock-96h
4hpi
24hpi
48hpi
72hpi
96hpi
CDS
3' UTR
5' UTR


Figure S2

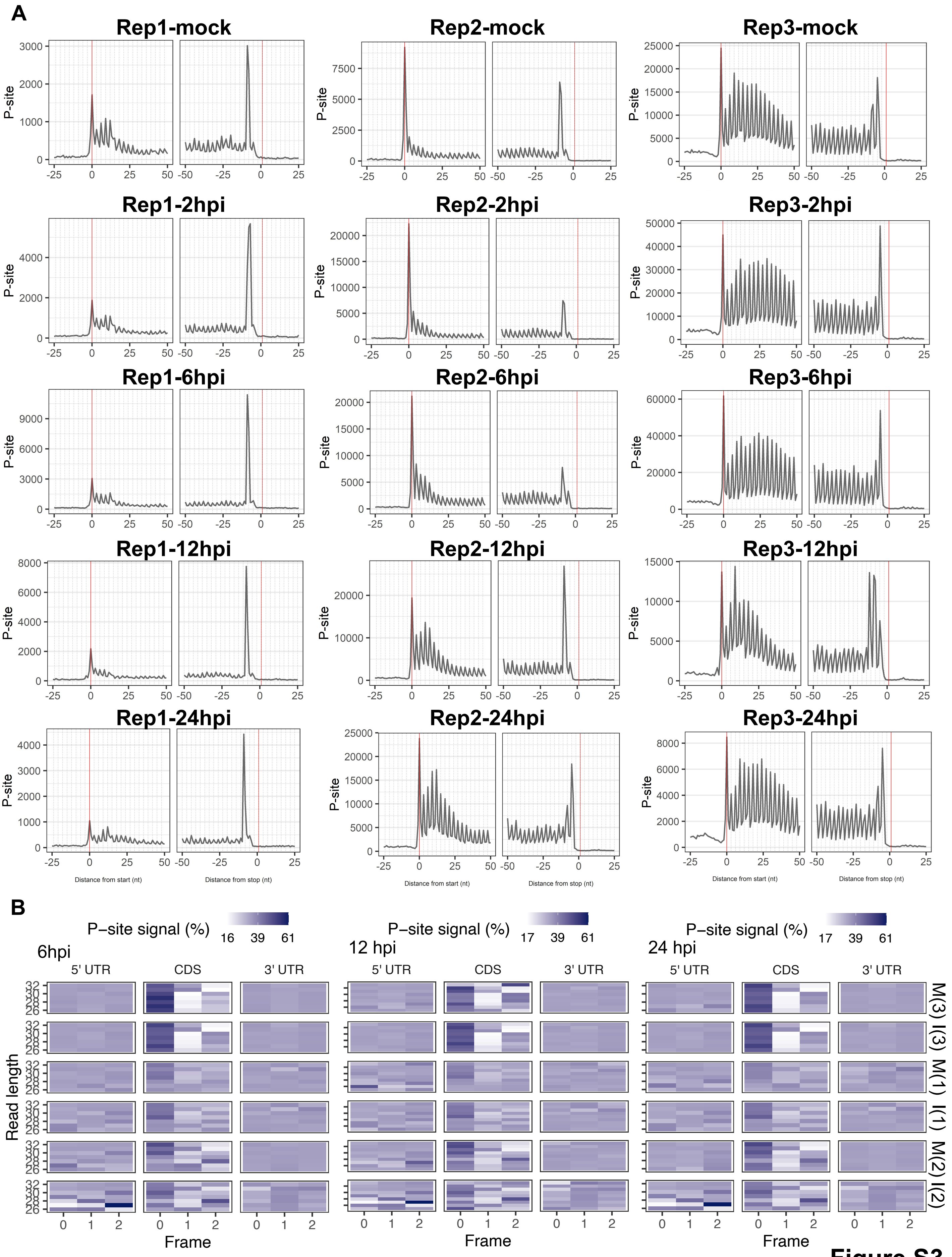
A
Rep1-mock
Rep2-mock
Rep3-mock
Rep1-2hpi
Rep2-2hpi
Rep3-2hpi
Rep1-6hpi
Rep2-6hpi
Rep3-6hpi
Rep1-12hpi
Rep2-12hpi
Rep3-12hpi
Rep1-24hpi
Rep2-24hpi
Rep3-24hpi
P-site
Distance from start (nt)
Distance from stop (nt)
B
P−site signal (%)
6hpi
12 hpi
24 hpi
5' UTR
CDS
3' UTR
Read length
Frame
M(3)
I(3)
M(1)
I(1)
M(2)
I(2)

Figure S3

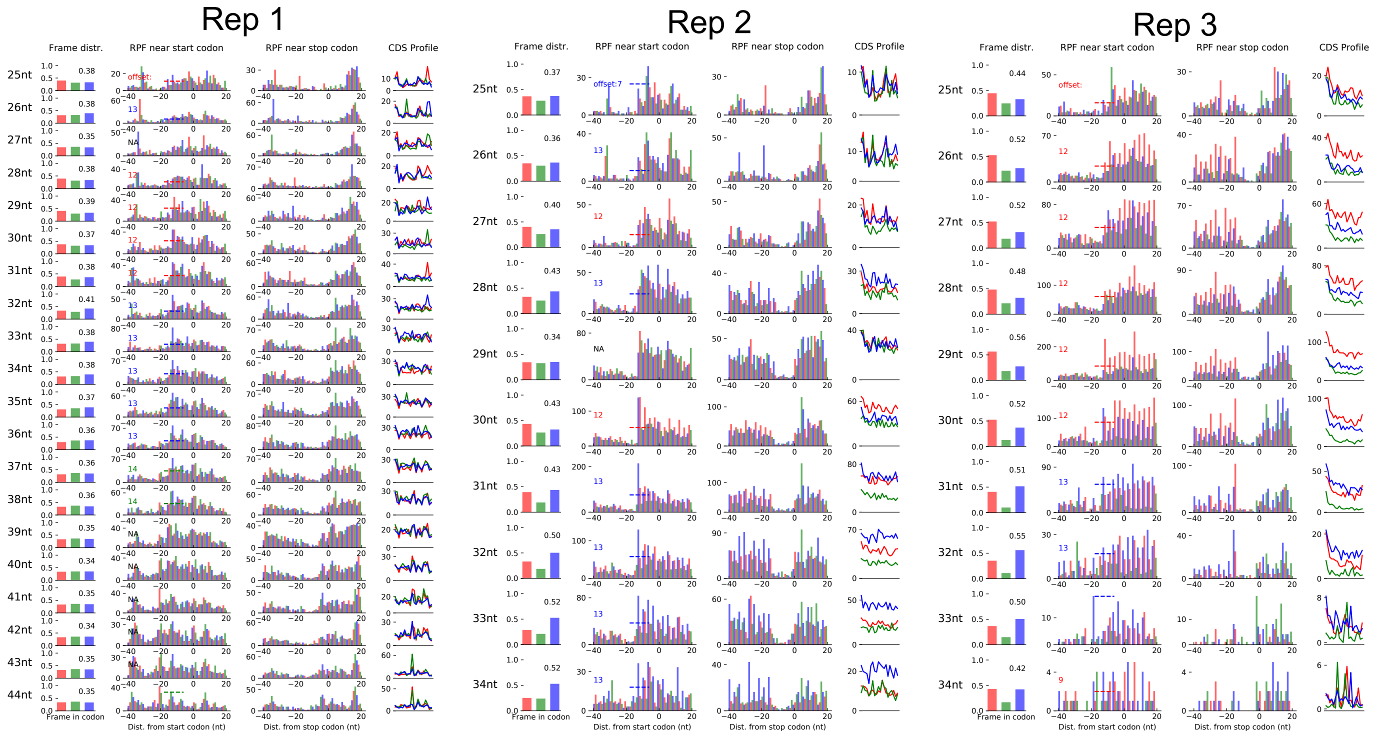

Rep 1
Rep 2
Rep 3
Frame distr.
RPF near start codon
RPF near stop codon
CDS Profile
Frame in codon
Dist. from start codon (nt)
Dist. from stop codon (nt)


**Figure S4**

**A**

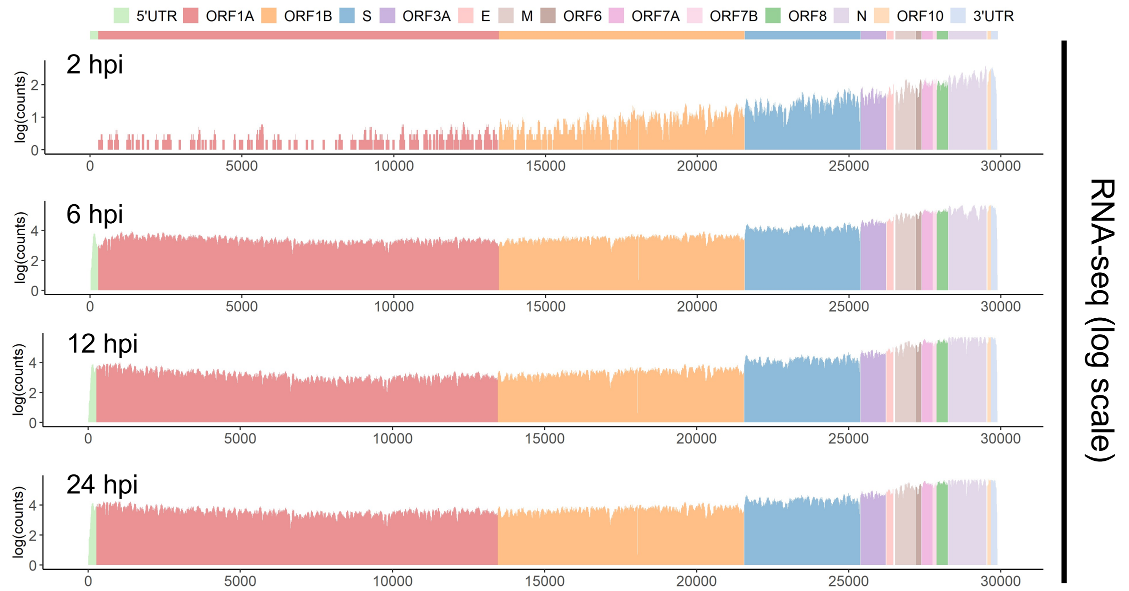
5'UTR ORF1A ORF1B S ORF3A E M ORF6 ORF7A ORF7B ORF8 N ORF10 3'UTR
2 hpi
6 hpi
12 hpi
24 hpi
log(counts)
0 5000 10000 15000 20000 25000 30000
RNA-seq (log scale)

**B**

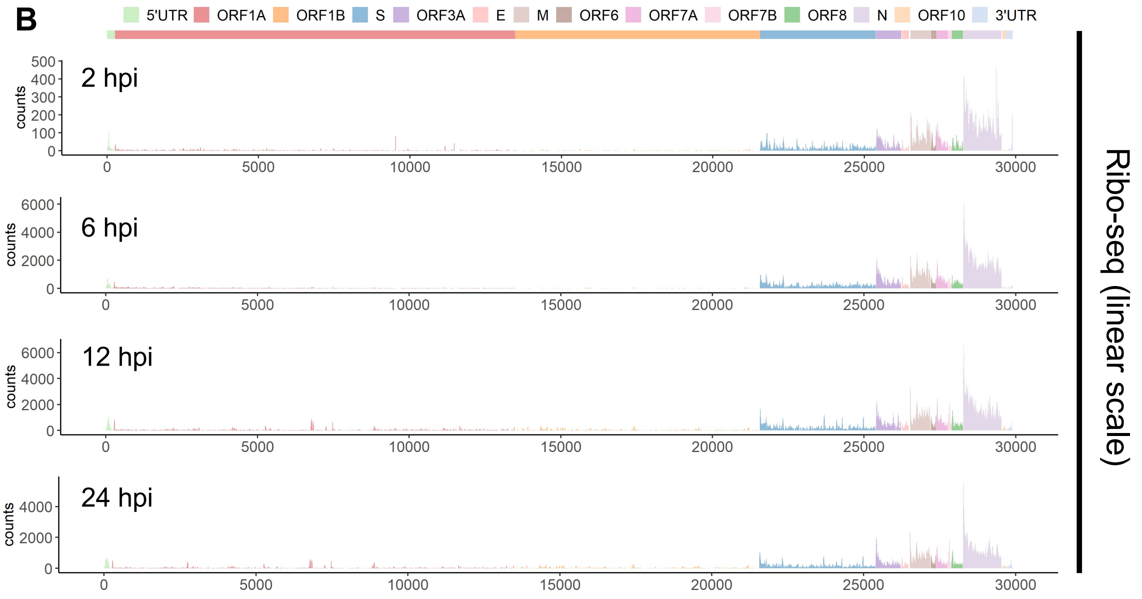
5'UTR ORF1A ORF1B S ORF3A E M ORF6 ORF7A ORF7B ORF8 N ORF10 3'UTR
2 hpi
6 hpi
12 hpi
24 hpi
counts
0 5000 10000 15000 20000 25000 30000
Ribo-seq (linear scale)

**Figure S5**

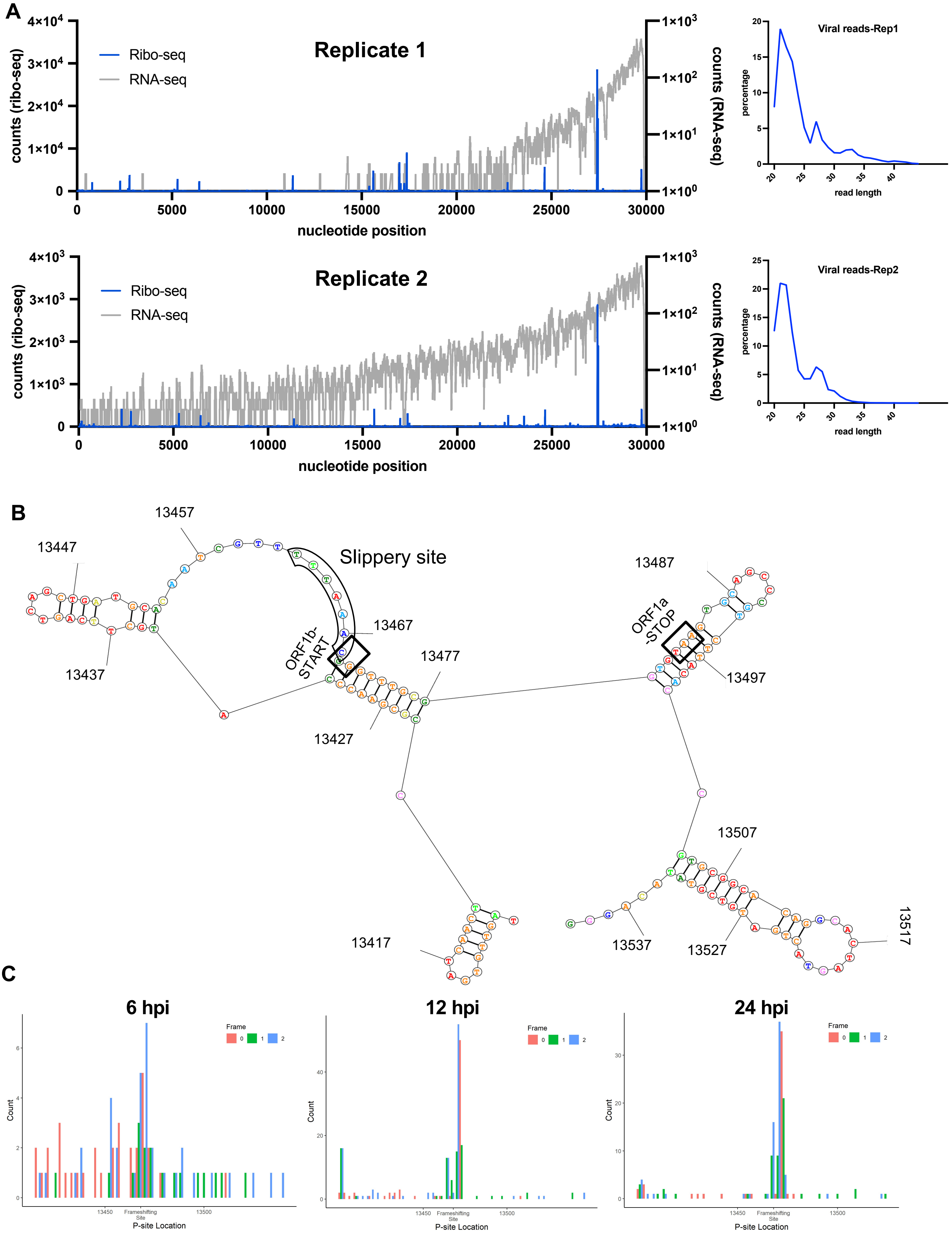
A
Replicate 1
Ribo-seq
RNA-seq
counts (ribo-seq)
counts (RNA-seq)
nucleotide position
Viral reads-Rep1
percentage
read length
Replicate 2
Viral reads-Rep2
B
Slippery site
ORF1b-START
ORF1a-STOP
13417
13427
13437
13447
13457
13467
13477
13487
13497
13507
13517
13527
13537
C
6 hpi
12 hpi
24 hpi
Frame
Count
P-site Location
Frameshifting Site

**Figure S6**

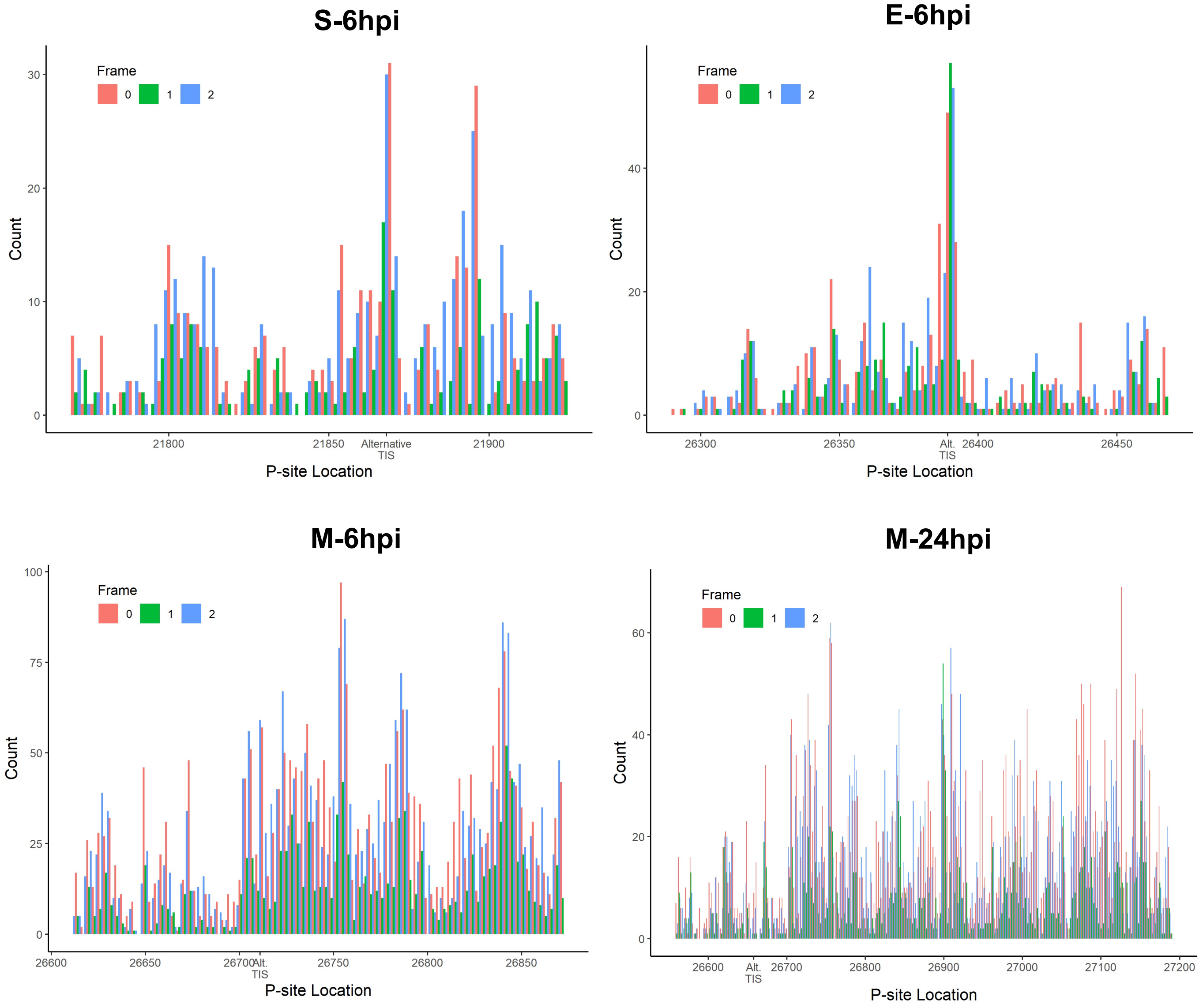
S-6hpi
Frame
0
1
2
Count
P-site Location
21800
21850
Alternative TIS
21900
E-6hpi
Alt. TIS
26300
26350
26400
26450
M-6hpi
26600
26650
26700
26750
26800
26850
M-24hpi
26900
27000
27100
27200

**Figure S7**

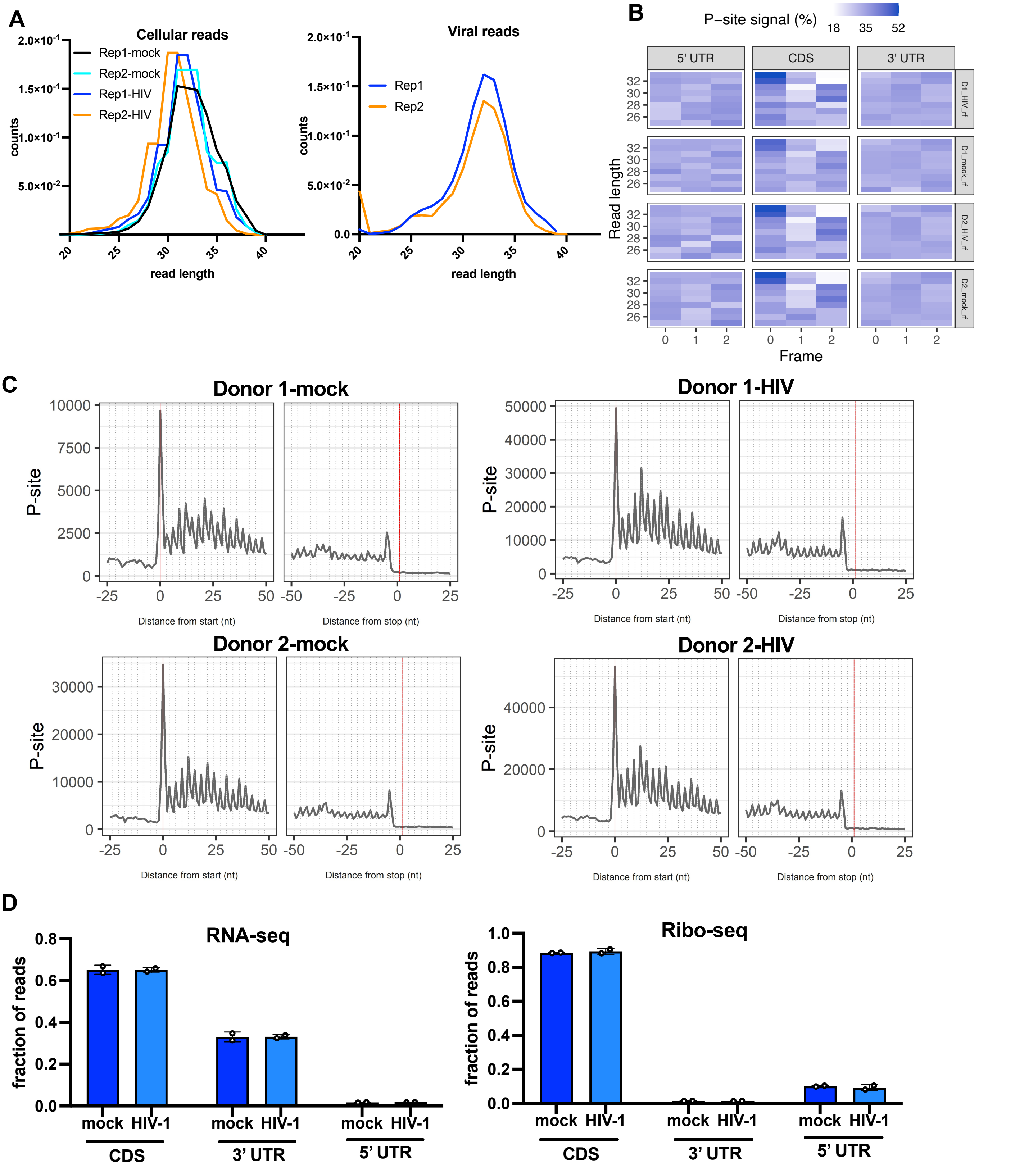
A
Cellular reads
Rep1-mock
Rep2-mock
Rep1-HIV
Rep2-HIV
counts
read length
Viral reads
Rep1
Rep2
B
P−site signal (%)
18
35
52
5' UTR
CDS
3' UTR
Read length
Frame
D1_HIV_rf
D1_mock_rf
D2_HIV_rf
D2_mock_rf
C
Donor 1-mock
Donor 1-HIV
Donor 2-mock
Donor 2-HIV
P-site
Distance from start (nt)
Distance from stop (nt)
D
RNA-seq
Ribo-seq
fraction of reads
mock
HIV-1
CDS
3' UTR
5' UTR

Figure S8

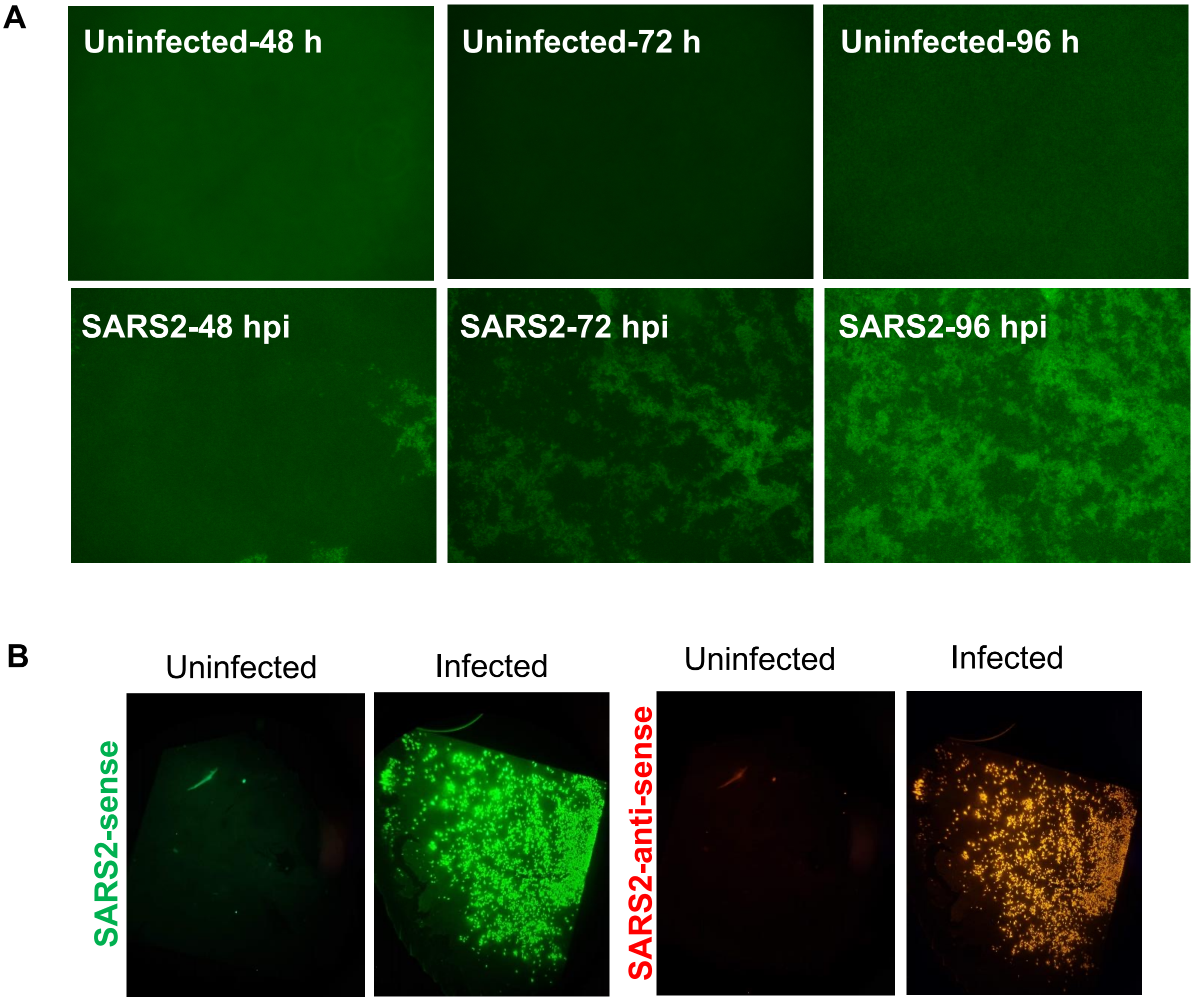
A
Uninfected-48 h
Uninfected-72 h
Uninfected-96 h
SARS2-48 hpi
SARS2-72 hpi
SARS2-96 hpi
B
Uninfected
Infected
Uninfected
Infected
SARS2-sense
SARS2-anti-sense

Figure S9

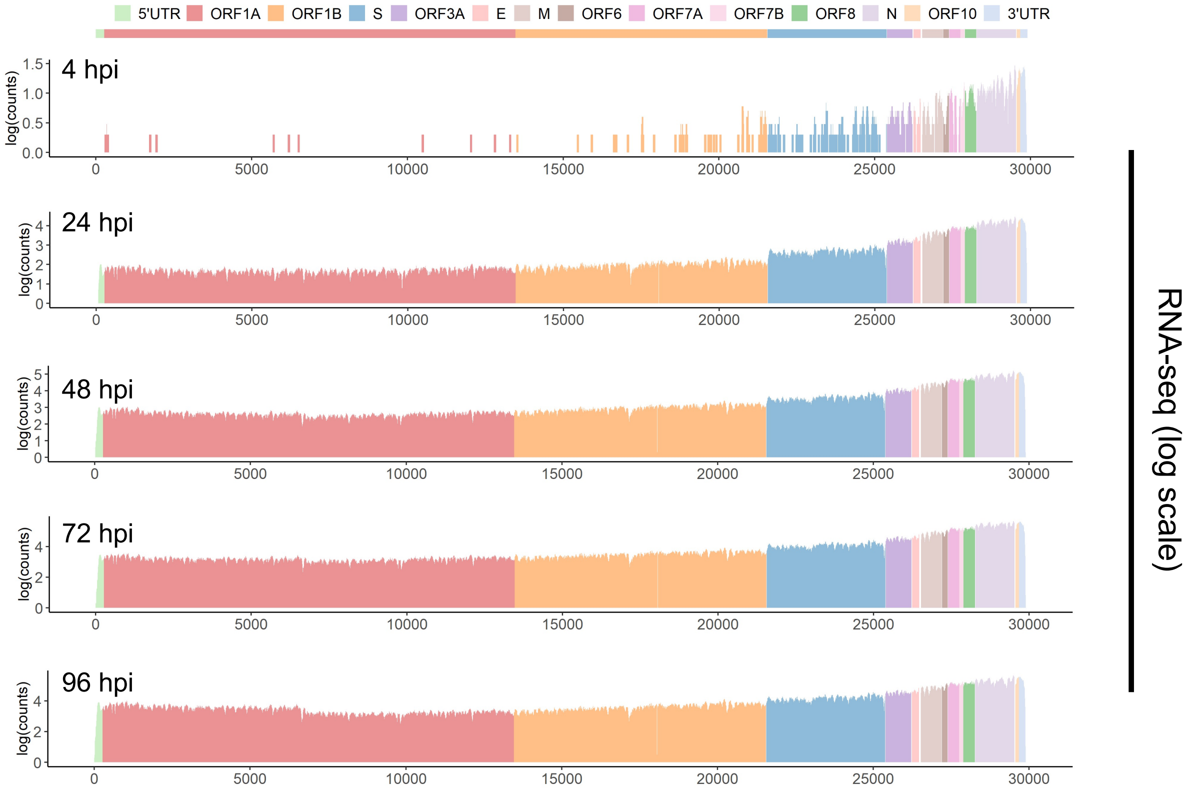
5'UTR
ORF1A
ORF1B
S
ORF3A
E
M
ORF6
ORF7A
ORF7B
ORF8
N
ORF10
3'UTR
4 hpi
24 hpi
48 hpi
72 hpi
96 hpi
log(counts)
RNA-seq (log scale)

**Figure S10**

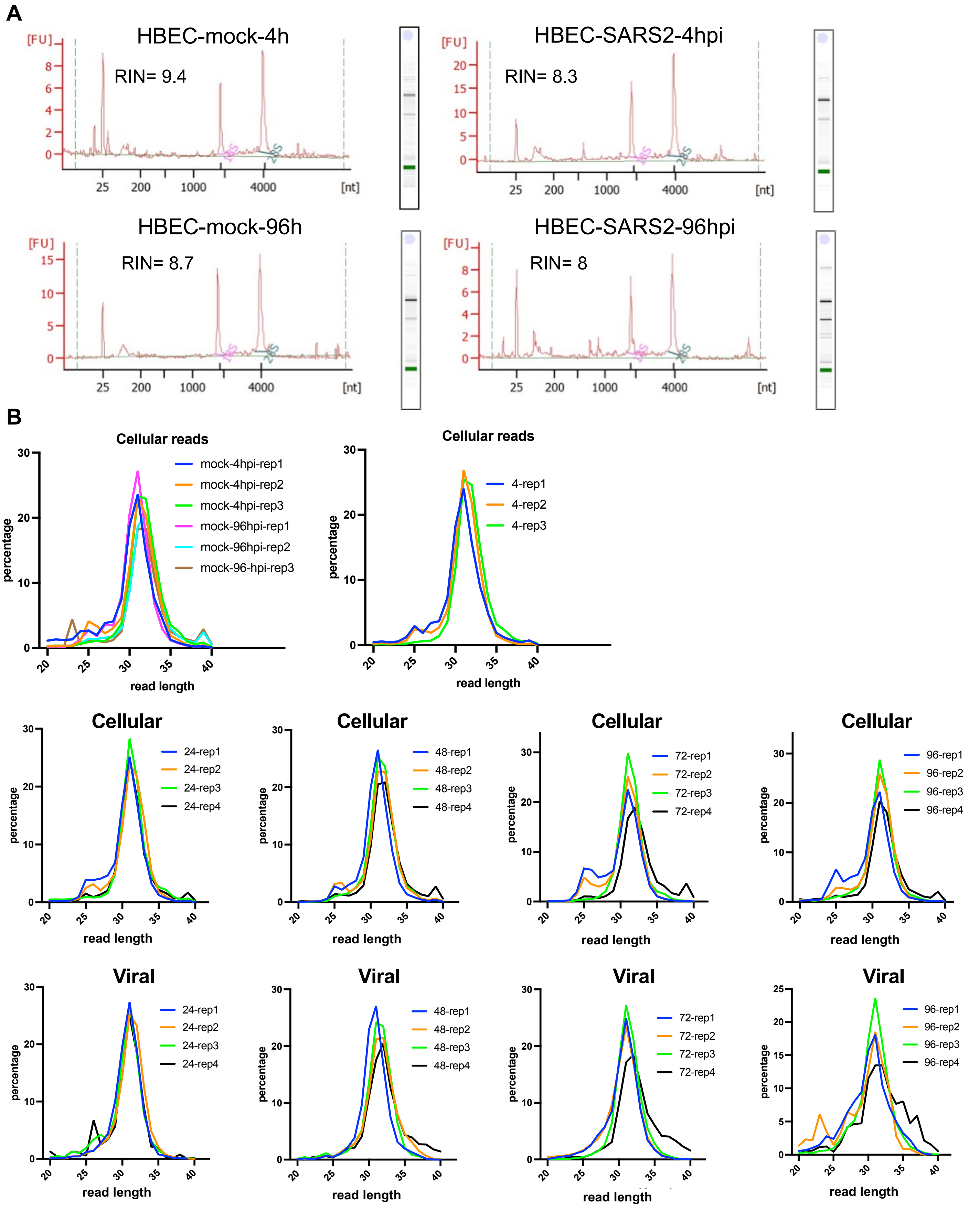
A
HBEC-mock-4h
RIN= 9.4
HBEC-SARS2-4hpi
RIN= 8.3
HBEC-mock-96h
RIN= 8.7
HBEC-SARS2-96hpi
RIN= 8
[FU]
18S
28S
25
200
1000
4000
[nt]
B
Cellular reads
mock-4hpi-rep1
mock-4hpi-rep2
mock-4hpi-rep3
mock-96hpi-rep1
mock-96hpi-rep2
mock-96-hpi-rep3
4-rep1
4-rep2
4-rep3
percentage
read length
Cellular
24-rep1
24-rep2
24-rep3
24-rep4
48-rep1
48-rep2
48-rep3
48-rep4
72-rep1
72-rep2
72-rep3
72-rep4
96-rep1
96-rep2
96-rep3
96-rep4
Viral

**Figure S11**

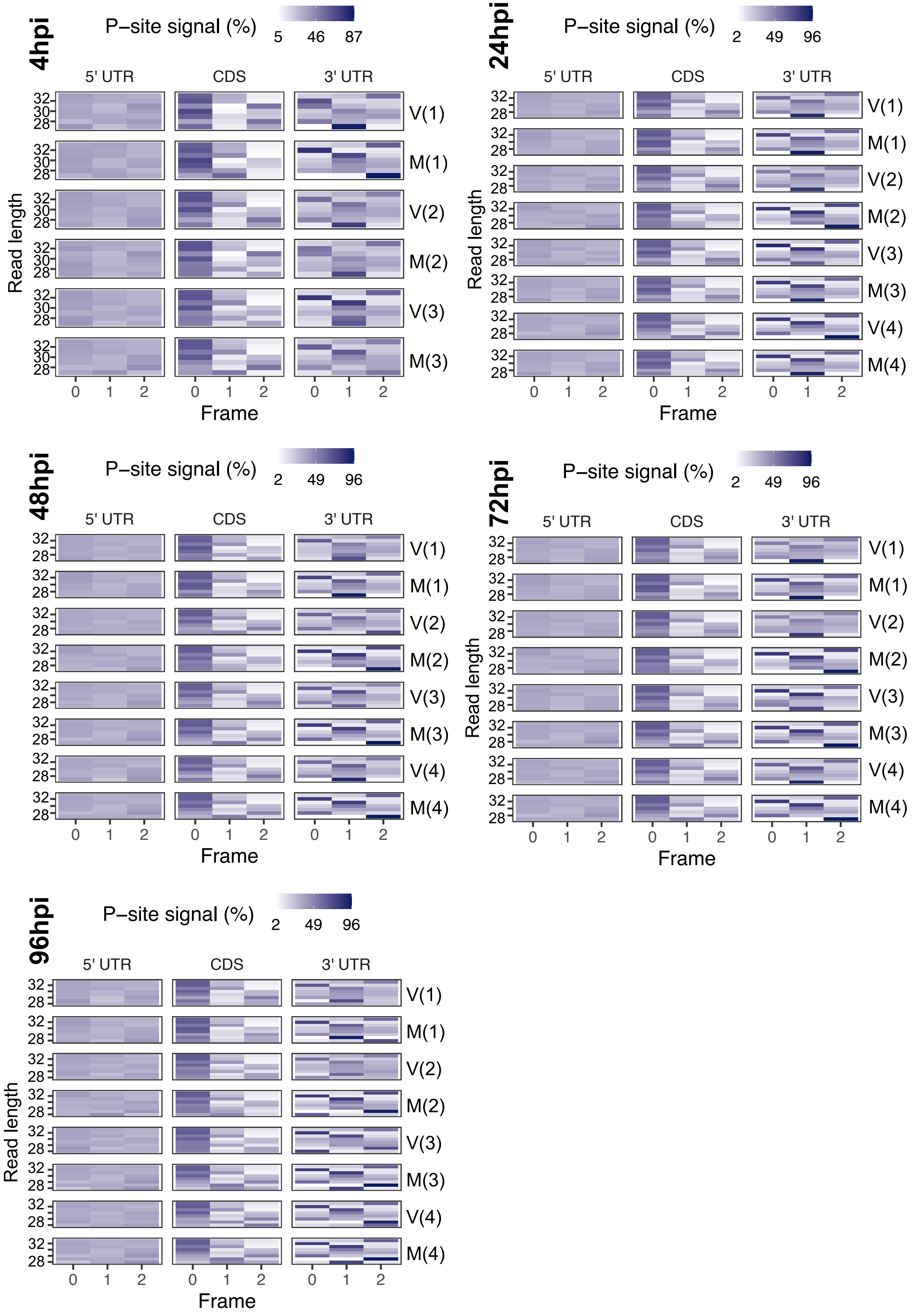
4hpi
P–site signal (%)
5 46 87
24hpi
P–site signal (%)
2 49 96
48hpi
P–site signal (%)
2 49 96
72hpi
P–site signal (%)
2 49 96
96hpi
P–site signal (%)
2 49 96
5' UTR
CDS
3' UTR
Read length
Frame
V(1)
M(1)
V(2)
M(2)
V(3)
M(3)
V(4)
M(4)

**Figure S12**

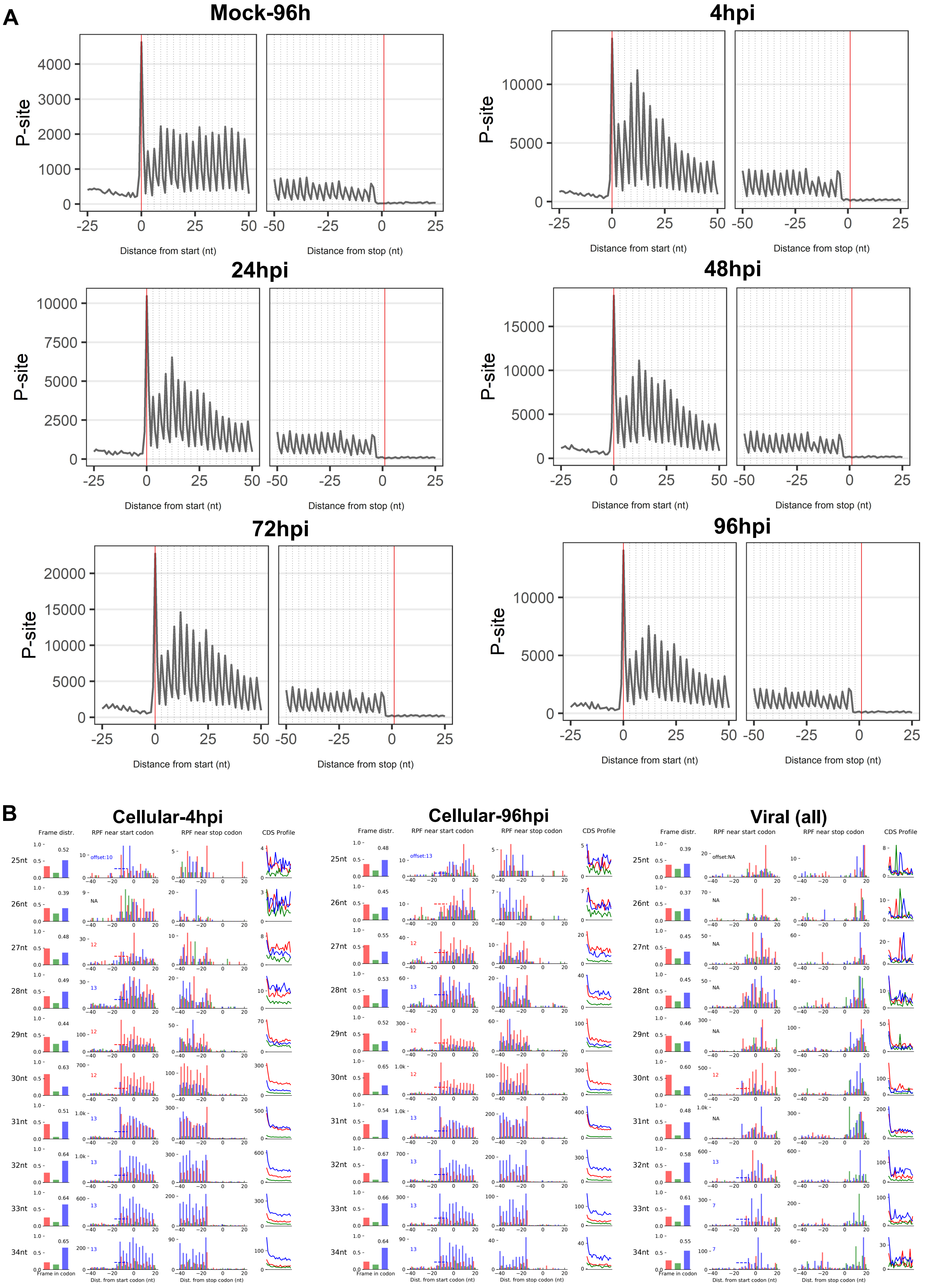
A
Mock-96h
4hpi
24hpi
48hpi
72hpi
96hpi
P-site
Distance from start (nt)
Distance from stop (nt)
B
Cellular-4hpi
Cellular-96hpi
Viral (all)
Frame distr.
RPF near start codon
RPF near stop codon
CDS Profile
25nt
26nt
27nt
28nt
29nt
30nt
31nt
32nt
33nt
34nt
Frame in codon
Dist. from start codon (nt)
Dist. from stop codon (nt)

Figure S13

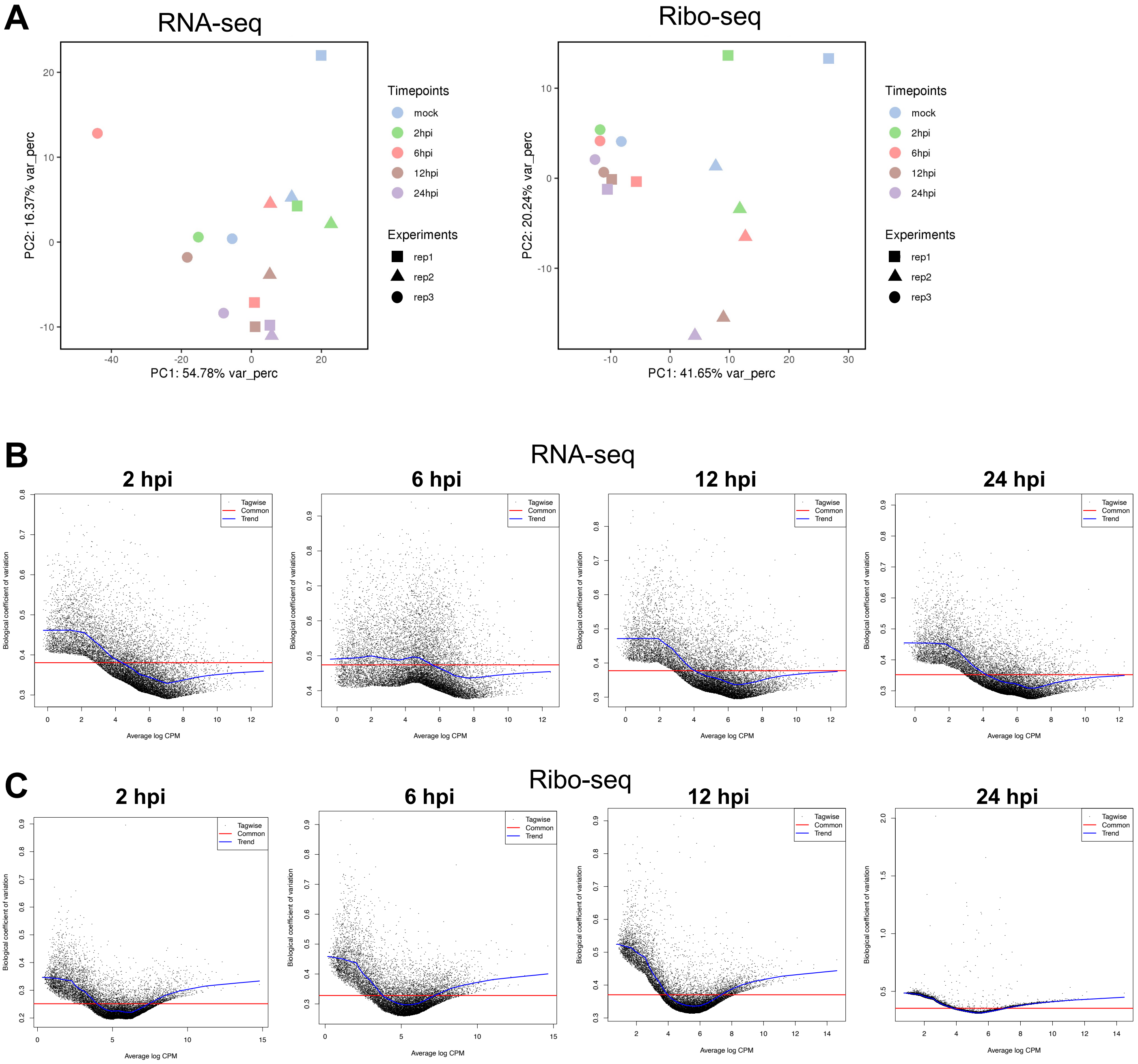
A
RNA-seq
Ribo-seq
PC2: 16.37% var_perc
PC1: 54.78% var_perc
PC2: 20.24% var_perc
PC1: 41.65% var_perc
Timepoints
mock
2hpi
6hpi
12hpi
24hpi
Experiments
rep1
rep2
rep3
B
RNA-seq
2 hpi
6 hpi
12 hpi
24 hpi
Tagwise
Common
Trend
Biological coefficient of variation
Average log CPM
C
Ribo-seq
2 hpi
6 hpi
12 hpi
24 hpi
Tagwise
Common
Trend
Biological coefficient of variation
Average log CPM

**Figure S14**

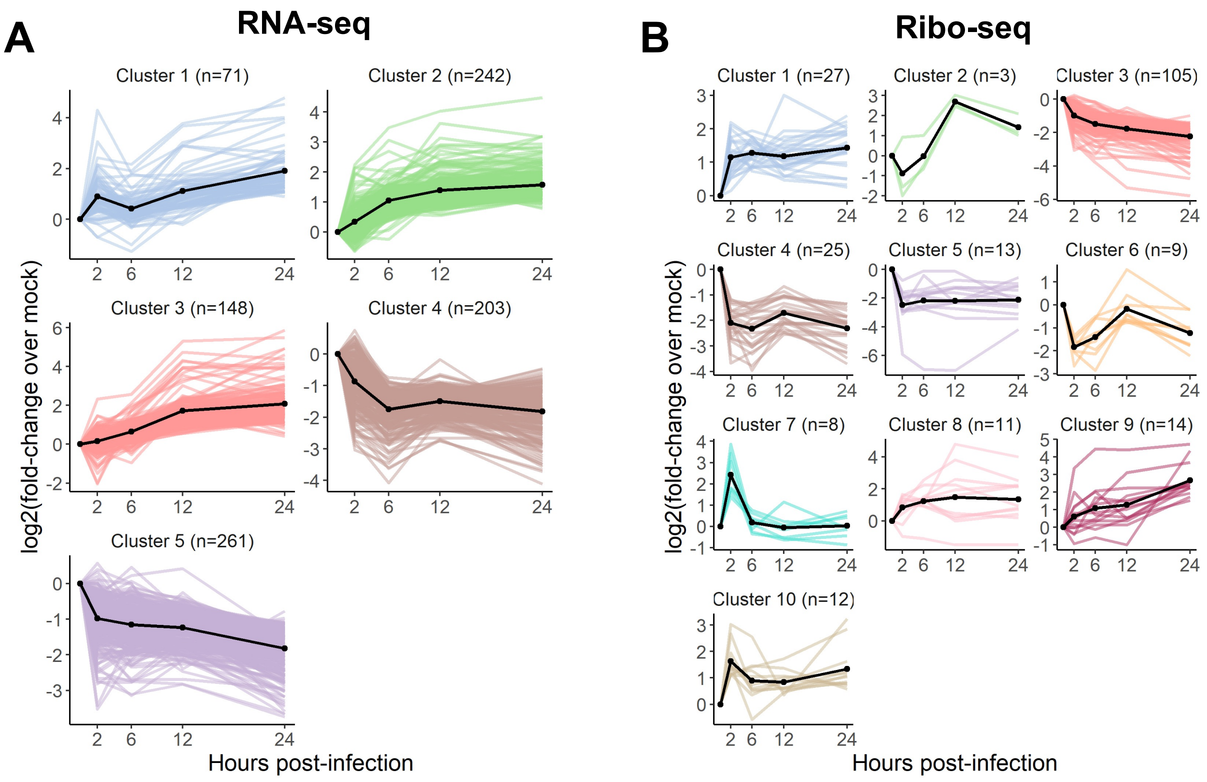
A
RNA-seq
Cluster 1 (n=71)
Cluster 2 (n=242)
Cluster 3 (n=148)
Cluster 4 (n=203)
Cluster 5 (n=261)
log2(fold-change over mock)
Hours post-infection
B
Ribo-seq
Cluster 1 (n=27)
Cluster 2 (n=3)
Cluster 3 (n=105)
Cluster 4 (n=25)
Cluster 5 (n=13)
Cluster 6 (n=9)
Cluster 7 (n=8)
Cluster 8 (n=11)
Cluster 9 (n=14)
Cluster 10 (n=12)
log2(fold-change over mock)
Hours post-infection

**Figure S15**

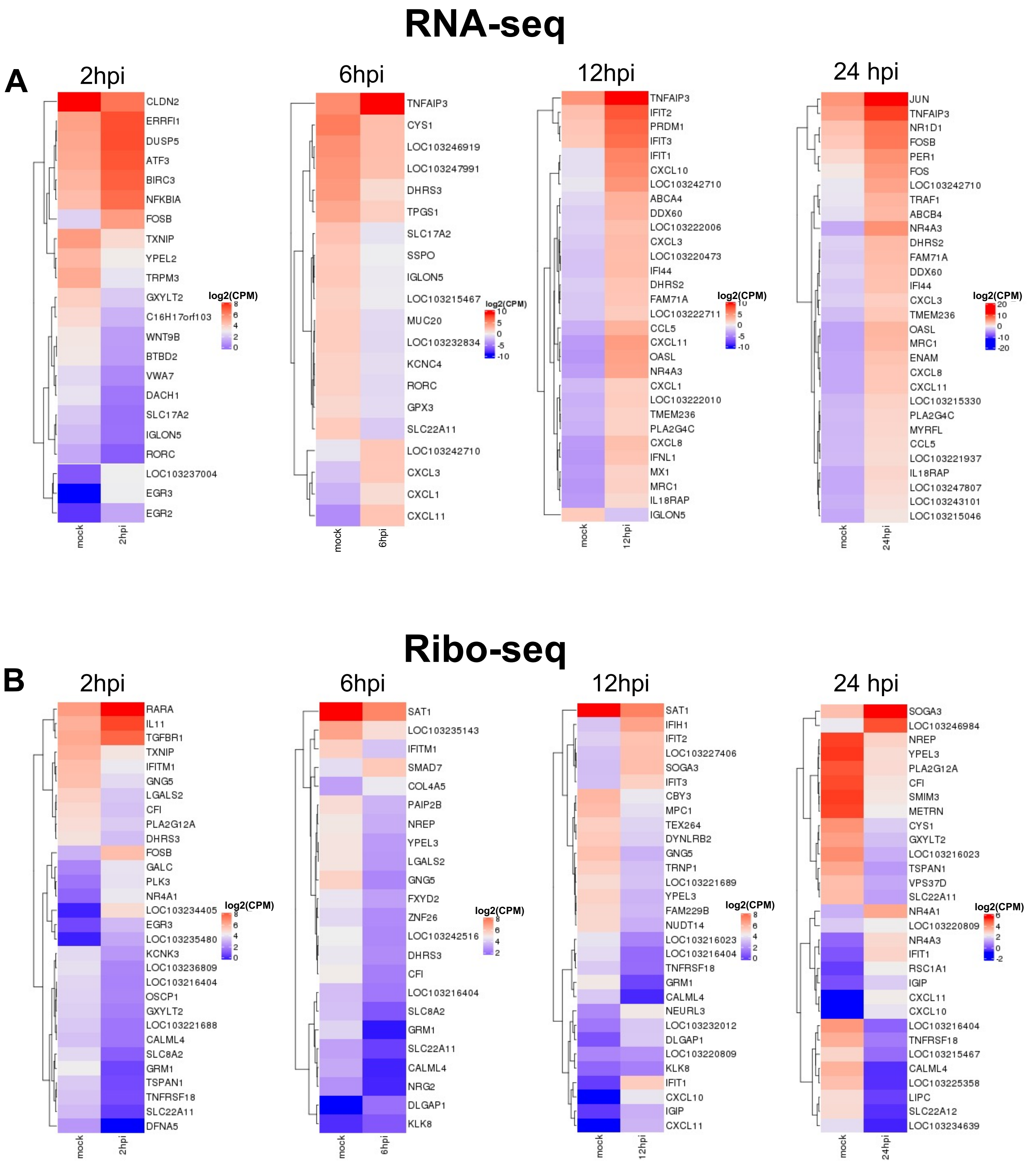
RNA-seq
A
2hpi
CLDN2
ERRFI1
DUSP5
ATF3
BIRC3
NFKBIA
FOSB
TXNIP
YPEL2
TRPM3
GXYLT2
C16H17orf103
WNT9B
BTBD2
VWA7
DACH1
SLC17A2
IGLON5
RORC
LOC103237004
EGR3
EGR2
mock
2hpi
log2(CPM)
6hpi
TNFAIP3
CYS1
LOC103246919
LOC103247991
DHRS3
TPGS1
SLC17A2
SSPO
IGLON5
LOC103215467
MUC20
LOC103232834
KCNC4
RORC
GPX3
SLC22A11
LOC103242710
CXCL3
CXCL1
CXCL11
mock
6hpi
log2(CPM)
12hpi
TNFAIP3
IFIT2
PRDM1
IFIT3
IFIT1
CXCL10
LOC103242710
ABCA4
DDX60
LOC103222006
CXCL3
LOC103220473
IFI44
DHRS2
FAM71A
LOC103222711
CCL5
CXCL11
OASL
NR4A3
CXCL1
LOC103222010
TMEM236
PLA2G4C
CXCL8
IFNL1
MX1
MRC1
IL18RAP
IGLON5
mock
12hpi
log2(CPM)
24 hpi
JUN
TNFAIP3
NR1D1
FOSB
PER1
FOS
LOC103242710
TRAF1
ABCB4
NR4A3
DHRS2
FAM71A
DDX60
IFI44
CXCL3
TMEM236
OASL
MRC1
ENAM
CXCL8
CXCL11
LOC103215330
PLA2G4C
MYRFL
CCL5
LOC103221937
IL18RAP
LOC103247807
LOC103243101
LOC103215046
mock
24hpi
log2(CPM)
Ribo-seq
B
2hpi
RARA
IL11
TGFBR1
TXNIP
IFITM1
GNG5
LGALS2
CFI
PLA2G12A
DHRS3
FOSB
GALC
PLK3
NR4A1
LOC103234405
EGR3
LOC103235480
KCNK3
LOC103236809
LOC103216404
OSCP1
GXYLT2
LOC103221688
CALML4
SLC8A2
GRM1
TSPAN1
TNFRSF18
SLC22A11
DFNA5
mock
2hpi
log2(CPM)
6hpi
SAT1
LOC103235143
IFITM1
SMAD7
COL4A5
PAIP2B
NREP
YPEL3
LGALS2
GNG5
FXYD2
ZNF26
LOC103242516
DHRS3
CFI
LOC103216404
SLC8A2
GRM1
SLC22A11
CALML4
NRG2
DLGAP1
KLK8
mock
6hpi
log2(CPM)
12hpi
SAT1
IFIH1
IFIT2
LOC103227406
SOGA3
IFIT3
CBY3
MPC1
TEX264
DYNLRB2
GNG5
TRNP1
LOC103221689
YPEL3
FAM229B
NUDT14
LOC103216023
LOC103216404
TNFRSF18
GRM1
CALML4
NEURL3
LOC103232012
DLGAP1
LOC103220809
KLK8
IFIT1
CXCL10
IGIP
CXCL11
mock
12hpi
log2(CPM)
24 hpi
SOGA3
LOC103246984
NREP
YPEL3
PLA2G12A
CFI
SMIM3
METRN
CYS1
GXYLT2
LOC103216023
TSPAN1
VPS37D
SLC22A11
NR4A1
LOC103220809
NR4A3
IFIT1
RSC1A1
IGIP
CXCL11
CXCL10
LOC103216404
TNFRSF18
LOC103215467
CALML4
LOC103225358
LIPC
SLC22A12
LOC103234639
mock
24hpi
log2(CPM)

Figure S16

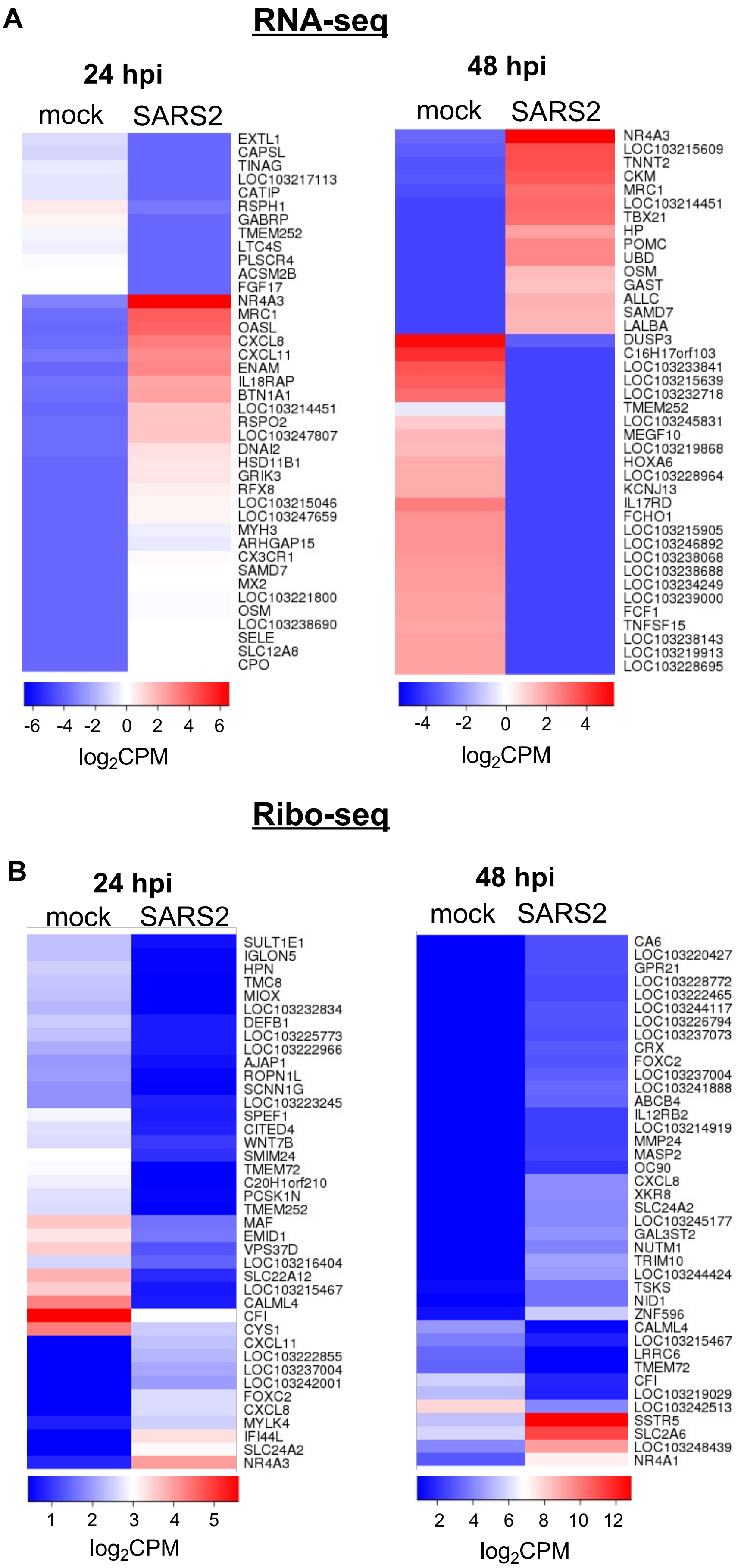
A
RNA-seq
24 hpi
mock
SARS2
EXTL1
CAPSL
TINAG
LOC103217113
CATIP
RSPH1
GABRP
TMEM252
LTC4S
PLSCR4
ACSM2B
FGF17
NR4A3
MRC1
OASL
CXCL8
CXCL11
ENAM
IL18RAP
BTN1A1
LOC103214451
RSPO2
LOC103247807
DNAI2
HSD11B1
GRIK3
RFX8
LOC103215046
LOC103247659
MYH3
ARHGAP15
CX3CR1
SAMD7
MX2
LOC103221800
OSM
LOC103238690
SELE
SLC12A8
CPO
-6 -4 -2 0 2 4 6
log2CPM
48 hpi
mock
SARS2
NR4A3
LOC103215609
TNNT2
CKM
MRC1
LOC103214451
TBX21
HP
POMC
UBD
OSM
GAST
ALLC
SAMD7
LALBA
DUSP3
C16H17orf103
LOC103233841
LOC103215639
LOC103232718
TMEM252
LOC103245831
MEGF10
LOC103219868
HOXA6
LOC103228964
KCNJ13
IL17RD
FCHO1
LOC103215905
LOC103246892
LOC103238068
LOC103238688
LOC103234249
LOC103239000
FCF1
TNFSF15
LOC103238143
LOC103219913
LOC103228695
-4 -2 0 2 4
log2CPM
Ribo-seq
B
24 hpi
mock
SARS2
SULT1E1
IGLON5
HPN
TMC8
MIOX
LOC103232834
DEFB1
LOC103225773
LOC103222966
AJAP1
ROPN1L
SCNN1G
LOC103223245
SPEF1
CITED4
WNT7B
SMIM24
TMEM72
C20H1orf210
PCSK1N
TMEM252
MAF
EMID1
VPS37D
LOC103216404
SLC22A12
LOC103215467
CALML4
CFI
CYS1
CXCL11
LOC103222855
LOC103237004
LOC103242001
FOXC2
CXCL8
MYLK4
IFI44L
SLC24A2
NR4A3
1 2 3 4 5
log2CPM
48 hpi
mock
SARS2
CA6
LOC103220427
GPR21
LOC103228772
LOC103222465
LOC103244117
LOC103226794
LOC103237073
CRX
FOXC2
LOC103237004
LOC103241888
ABCB4
IL12RB2
LOC103214919
MMP24
MASP2
OC90
CXCL8
XKR8
SLC24A2
LOC103245177
GAL3ST2
NUTM1
TRIM10
LOC103244424
TSKS
NID1
ZNF596
CALML4
LOC103215467
LRRC6
TMEM72
CFI
LOC103219029
LOC103242513
SSTR5
SLC2A6
LOC103248439
NR4A1
2 4 6 8 10 12
log2CPM

**Figure S17**

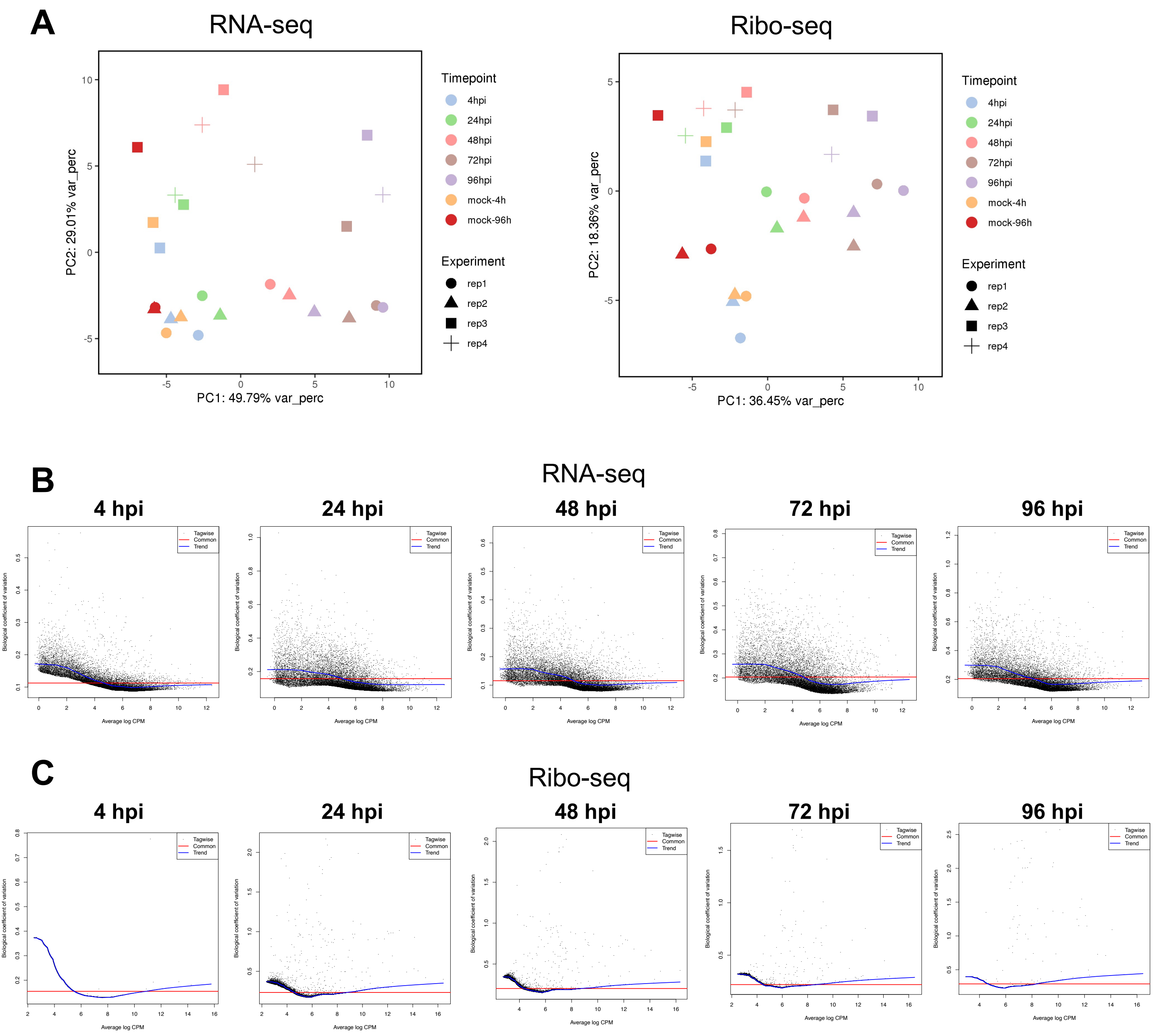

A
RNA-seq
PC1: 49.79% var_perc
PC2: 29.01% var_perc
Ribo-seq
PC1: 36.45% var_perc
PC2: 18.36% var_perc
Timepoint
4hpi
24hpi
48hpi
72hpi
96hpi
mock-4h
mock-96h
Experiment
rep1
rep2
rep3
rep4
B
RNA-seq
4 hpi
24 hpi
48 hpi
72 hpi
96 hpi
Tagwise
Common
Trend
Biological coefficient of variation
Average log CPM
C
Ribo-seq
4 hpi
24 hpi
48 hpi
72 hpi
96 hpi
Tagwise
Common
Trend
Biological coefficient of variation
Average log CPM


**Figure S18**

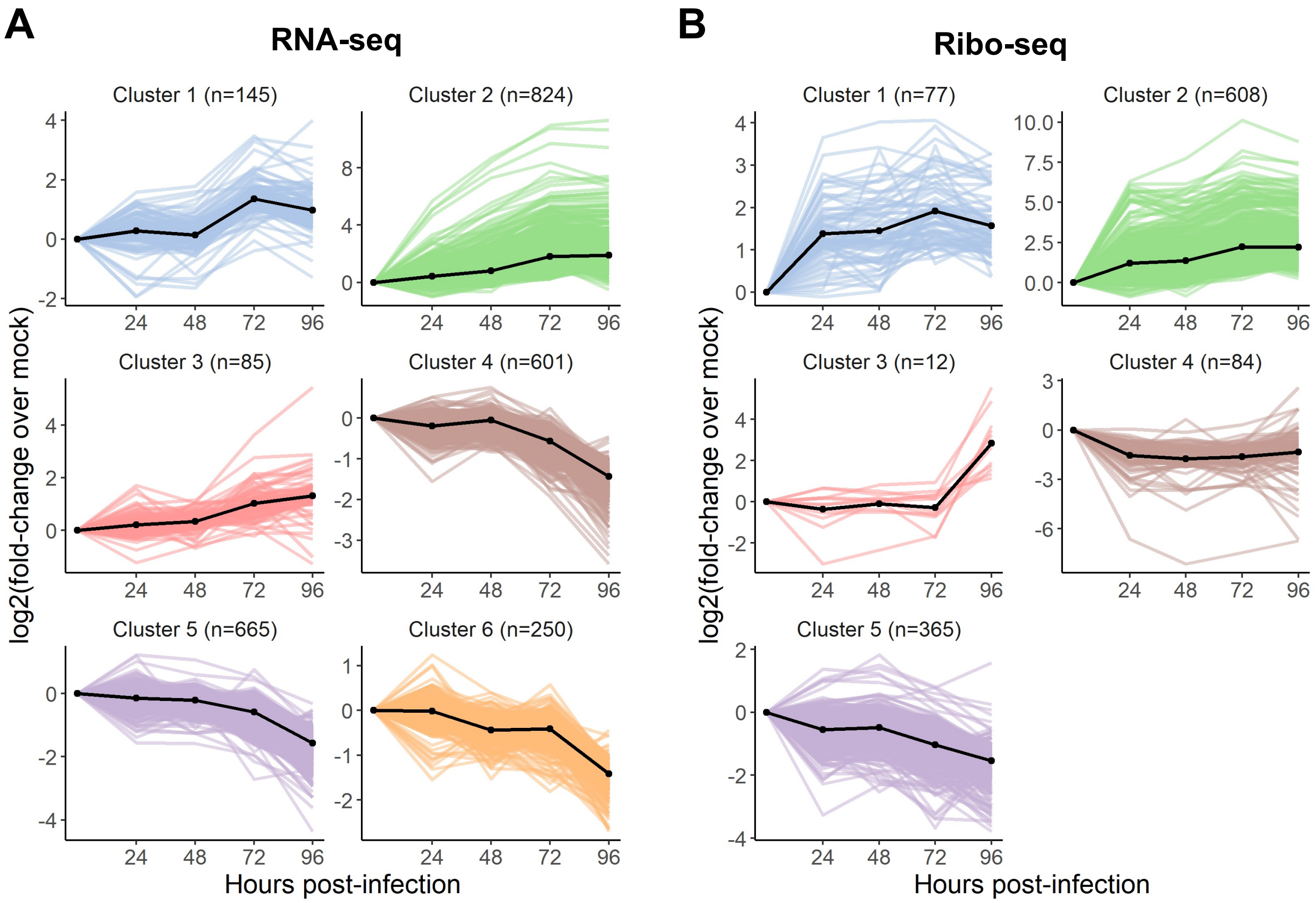
A
RNA-seq
Cluster 1 (n=145)
Cluster 2 (n=824)
Cluster 3 (n=85)
Cluster 4 (n=601)
Cluster 5 (n=665)
Cluster 6 (n=250)
log2(fold-change over mock)
Hours post-infection
B
Ribo-seq
Cluster 1 (n=77)
Cluster 2 (n=608)
Cluster 3 (n=12)
Cluster 4 (n=84)
Cluster 5 (n=365)
log2(fold-change over mock)
Hours post-infection

Figure S19

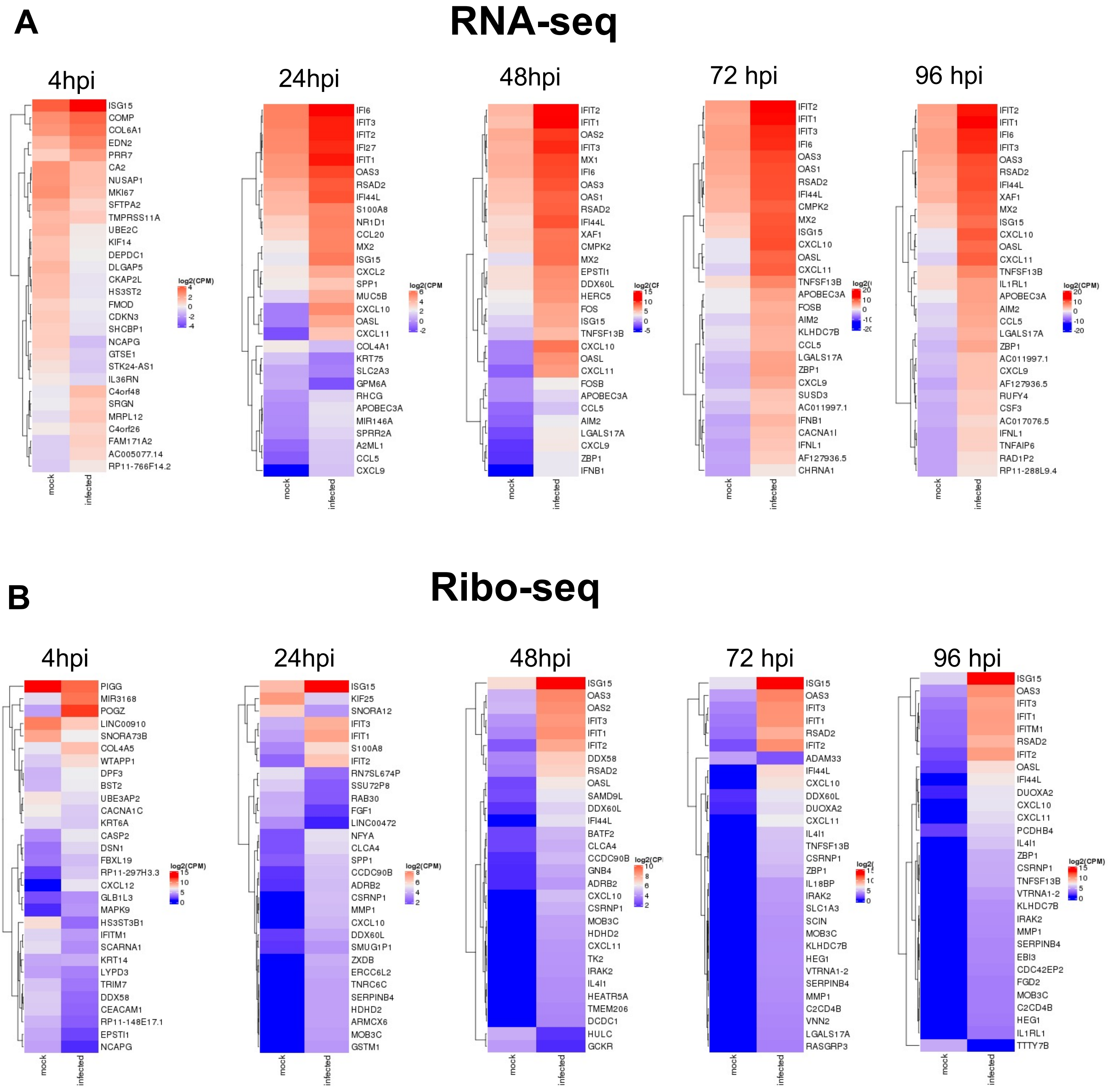
A
RNA-seq
4hpi
ISG15 COMP COL6A1 EDN2 PRR7 CA2 NUSAP1 MKI67 SFTPA2 TMPRSS11A UBE2C KIF14 DEPDC1 DLGAP5 CKAP2L HS3ST2 FMOD CDKN3 SHCBP1 NCAPG GTSE1 STK24-AS1 IL36RN C4orf48 SRGN MRPL12 C4orf26 FAM171A2 AC005077.14 RP11-766F14.2
log2(CPM)
mock
infected
24hpi
IFI6 IFIT3 IFIT2 IFI27 IFIT1 OAS3 RSAD2 IFI44L S100A8 NR1D1 CCL20 MX2 ISG15 CXCL2 SPP1 MUC5B CXCL10 OASL CXCL11 COL4A1 KRT75 SLC2A3 GPM6A RHCG APOBEC3A MIR146A SPRR2A A2ML1 CCL5 CXCL9
mock
infected
48hpi
IFIT2 IFIT1 OAS2 IFIT3 MX1 IFI6 OAS3 OAS1 RSAD2 IFI44L XAF1 CMPK2 MX2 EPSTI1 DDX60L HERC5 FOS ISG15 TNFSF13B CXCL10 OASL CXCL11 FOSB APOBEC3A CCL5 AIM2 LGALS17A CXCL9 ZBP1 IFNB1
mock
infected
72 hpi
IFIT2 IFIT1 IFIT3 IFI6 OAS3 OAS1 RSAD2 IFI44L CMPK2 MX2 ISG15 CXCL10 OASL CXCL11 TNFSF13B APOBEC3A FOSB AIM2 KLHDC7B CCL5 LGALS17A ZBP1 CXCL9 SUSD3 AC011997.1 IFNB1 CACNA1I IFNL1 AF127936.5 CHRNA1
mock
infected
96 hpi
IFIT2 IFIT1 IFI6 IFIT3 OAS3 RSAD2 IFI44L XAF1 MX2 ISG15 CXCL10 OASL CXCL11 TNFSF13B IL1RL1 APOBEC3A AIM2 CCL5 LGALS17A ZBP1 AC011997.1 CXCL9 AF127936.5 RUFY4 CSF3 AC017076.5 IFNL1 TNFAIP6 RAD1P2 RP11-288L9.4
log2(CPM)
mock
infected
B
Ribo-seq
4hpi
PIGG MIR3168 POGZ LINC00910 SNORA73B COL4A5 WTAPP1 DPF3 BST2 UBE3AP2 CACNA1C KRT6A CASP2 DSN1 FBXL19 RP11-297H3.3 CXCL12 GLB1L3 MAPK9 HS3ST3B1 IFITM1 SCARNA1 KRT14 LYPD3 TRIM7 DDX58 CEACAM1 RP11-148E17.1 EPSTI1 NCAPG
log2(CPM)
mock
infected
24hpi
ISG15 KIF25 SNORA12 IFIT3 IFIT1 S100A8 IFIT2 RN7SL674P SSU72P8 RAB30 FGF1 LINC00472 NFYA CLCA4 SPP1 CCDC90B ADRB2 CSRNP1 MMP1 CXCL10 DDX60L SMUG1P1 ZXDB ERCC6L2 TNRC6C SERPINB4 HDHD2 ARMCX6 MOB3C GSTM1
log2(CPM)
mock
infected
48hpi
ISG15 OAS3 OAS2 IFIT3 IFIT1 IFIT2 DDX58 RSAD2 OASL SAMD9L DDX60L IFI44L BATF2 CLCA4 CCDC90B GNB4 ADRB2 CXCL10 CSRNP1 MOB3C HDHD2 CXCL11 TK2 IRAK2 IL4I1 HEATR5A TMEM206 DCDC1 HULC GCKR
mock
infected
72 hpi
ISG15 OAS3 IFIT3 IFIT1 RSAD2 IFIT2 ADAM33 IFI44L CXCL10 DDX60L DUOXA2 CXCL11 IL4I1 TNFSF13B CSRNP1 ZBP1 IL18BP IRAK2 SLC1A3 SCIN MOB3C KLHDC7B HEG1 VTRNA1-2 SERPINB4 MMP1 C2CD4B VNN2 LGALS17A RASGRP3
mock
infected
96 hpi
ISG15 OAS3 IFIT3 IFIT1 IFITM1 RSAD2 IFIT2 OASL IFI44L DUOXA2 CXCL10 CXCL11 PCDHB4 IL4I1 ZBP1 CSRNP1 TNFSF13B VTRNA1-2 KLHDC7B IRAK2 MMP1 SERPINB4 EBI3 CDC42EP2 FGD2 MOB3C C2CD4B HEG1 IL1RL1 TTTY7B
log2(CPM)
mock
infected

**Figure S20**
